# Crowdsourcing assessment of maternal blood multi-omics for predicting gestational age and preterm birth

**DOI:** 10.1101/2020.06.05.130971

**Authors:** Adi L. Tarca, Bálint Ármin Pataki, Roberto Romero, Marina Sirota, Yuanfang Guan, Rintu Kutum, Nardhy Gomez-Lopez, Bogdan Done, Gaurav Bhatti, Thomas Yu, Gaia Andreoletti, Tinnakorn Chaiworapongsa, The DREAM Preterm Birth Prediction Challenge Consortium, Sonia S. Hassan, Chaur-Dong Hsu, Nima Aghaeepour, Gustavo Stolovitzky, Istvan Csabai, James C. Costello

## Abstract

Identification of pregnancies at risk of preterm birth (PTB), the leading cause of newborn deaths, remains challenging given the syndromic nature of the disease. We report a longitudinal multi-omics study coupled with a DREAM challenge to develop predictive models of PTB. We found that whole blood gene expression predicts ultrasound-based gestational ages in normal and complicated pregnancies (r=0.83), as well as the delivery date in normal pregnancies (r=0.86), with an accuracy comparable to ultrasound. However, unlike the latter, transcriptomic data collected at <37 weeks of gestation predicted the delivery date of one third of spontaneous (sPTB) cases within 2 weeks of the actual date. Based on samples collected before 33 weeks in asymptomatic women we found expression changes preceding preterm prelabor rupture of the membranes that were consistent across time points and cohorts, involving, among others, leukocyte-mediated immunity. Plasma proteomic random forests predicted sPTB with higher accuracy and earlier in pregnancy than whole blood transcriptomic models (e.g. AUROC=0.76 vs. AUROC=0.6 at 27-33 weeks of gestation).

## Introduction

Early identification of patients at risk for obstetrical disease is required to improve health outcomes and develop new therapeutic interventions. One of the “great obstetrical syndromes”^1^, preterm birth, defined as birth prior to the completion of 37 weeks of gestation, is the leading cause of newborn deaths worldwide. In 2010, 14.9 million babies were born preterm, accounting for 11.1% of all births across 184 countries—the highest preterm birth rates occurring in Africa and North America^2^. In the United States, the rate of prematurity remained fundamentally unchanged in recent years^3^ and it has an annual societal economic burden of at least $26.2 billion^4^. The high incidence of preterm birth is concerning: 29% of all neonatal deaths worldwide, approximately 1 million deaths in total, can be attributed to complications of prematurity^5^. Furthermore, children born prematurely are at increased risk for several short- and long-term complications that may include motor, cognitive, and behavioral impairments^6,7^.

Approximately one-third of preterm births are medically indicated for maternal (e.g. preeclampsia) or fetal conditions (e.g. growth restriction); the other two-thirds are categorized as spontaneous preterm births, inclusive of spontaneous preterm labor and delivery with intact membranes (sPTD), and preterm prelabor rupture of the membranes (PPROM)^8^. Preterm birth is a syndrome with multiple etiologies^9^, and its complexity makes accurate prediction by a single set of biomarkers difficult. While genetic risk factors for preterm birth have been reported^10^, the two most powerful predictors of spontaneous preterm birth are a sonographic short cervix in the midtrimester, and a history of spontaneous preterm birth in a prior pregnancy.^11^ As for prevention of the syndrome, vaginal progesterone administered to asymptomatic women with a short cervix in the midtrimester reduces the rate of preterm birth < 33 weeks of gestation by 45% and decreases the rate of neonatal complications, including neonatal respiratory distress syndrome^12–14^. The role of 17-alpha-hydroxyprogesterone caproate for preventing preterm delivery in patients with an episode of preterm labor is controversial^15,16^.

To compensate for the suboptimal prediction of preterm birth by currently used biomarkers, alternative approaches to identify biomarkers have been proposed, such as focusing on fetal and placenta-specific signatures^17^, with the latter eventually refined by single-cell genomics^17,18^, and by expanding the types of data collected via multi-omics platforms^10,19,20^. While molecular profiles have been shown to be strongly modulated by advancing gestation in the maternal blood proteome^21,22^, transcriptome^17,23^, and vaginal microbiome^24,25^, the timing of delivery based on such molecular clocks of pregnancy is still challenging^17^. A recent meta-analysis^26^ suggests that specific changes in the maternal whole blood transcriptome associated with spontaneous preterm birth are largely consistent across studies when both symptomatic and asymptomatic cases are involved and when the samples collected at or near the time of preterm delivery are also included. However, the accuracy of predictive models to make inferences in asymptomatic women early in pregnancy has not been evaluated. This topic is important, since early identification is necessary to develop treatment strategies to reduce the impact of prematurity.

Therefore, we generated longitudinal whole blood transcriptomic and plasma proteomic data on 216 women and leveraged the Dialogue for Reverse Engineering Assessments and Methods (DREAM) crowdsourcing framework^27^ to engage over 500 members of the computational biology community and robustly assess the value of maternal blood multi-omics data in two sub-challenges. In sub-challenge 1, we assessed maternal whole blood transcriptomic data for prediction of gestational age in normal and complicated pregnancies using the last menstrual period (LMP) and ultrasound estimate as the gold standard, and showed that predictions are robust to disease-related perturbations. To avoid potential biases in the gold standard, in a post-challenge analysis, we have also predicted delivery dates in women with spontaneous birth (**Fig. 1**), and found similar prediction performance. In sub-challenge 2, we evaluated within- and cross-cohort prediction of preterm birth leveraging longitudinal transcriptomic data in asymptomatic women generated herein and by Heng et al.^28^ in a cohort in Calgary. The separate consideration of both spontaneous preterm birth phenotypes, i.e. spontaneous preterm delivery with intact membranes (sPTD) and premature prelabor rupture of membranes (PPROM), allowed us to pinpoint that previously reported leukocyte activation-related RNA changes preceding preterm birth are shared across cohorts only for the PPROM phenotype but not sPTD. Moreover, the evaluation of plasma proteomics and blood multi-omics data to determine the earliest stage in gestation when biomarkers have predictive value (**Fig. 1**), also make this study unique, and led to the conclusion that changes in plasma proteomics can be detected earlier and are more accurate than whole blood transcripomics for prediction of preterm birth. In addition to the transcriptomic signatures of gestational age and the multi-omics signatures of preterm birth that were identified herein, this work sets a benchmark for evaluation of longitudinal omics data in pregnancy research. The computational lessons and algorithms for risk prediction from longitudinal omics data derived herein can also impact future studies.

**Figure 1.**
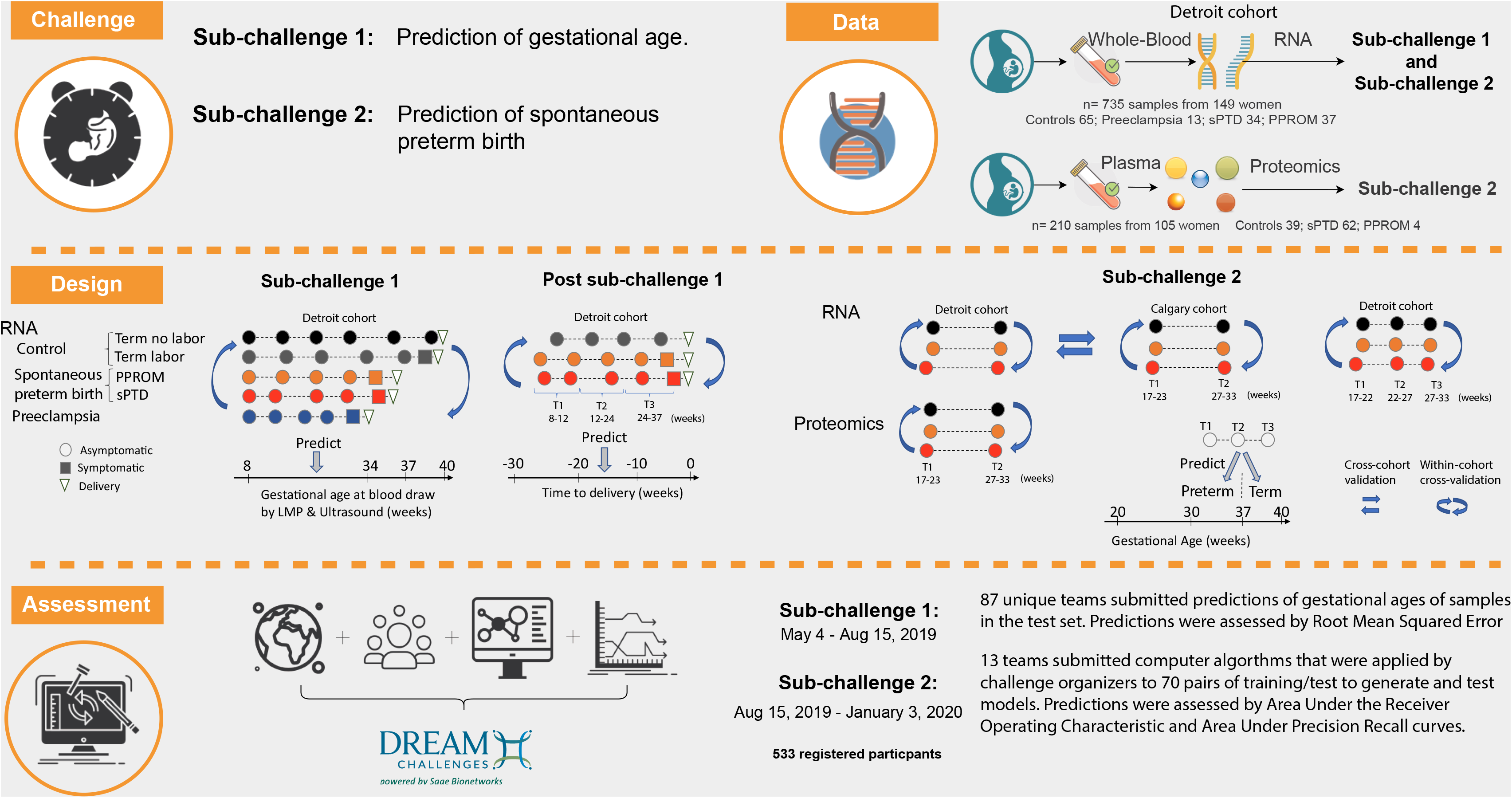
Study overview. Whole blood transcriptomic and/or plasma proteomic profiles were generated from 216 women with either normal pregnancy, spontaneous preterm birth with intact (sPTD) or ruptured membranes (PPROM), or preeclampsia. **Sub-challenge 1**: Whole blood transcriptomic data were generated from samples collected in normal pregnancies and those complicated by spontaneous preterm birth with intact (sPTD) or ruptured membranes (PPROM), or preeclampsia. Participating teams were provided gene expression data to develop their own prediction models for gestational age at blood draw defined by last menstrual period (LMP) and ultrasound (gold standard). Participants submitted predictions on a blinded test set (see **Fig. S1** for training/test partition). In a post challenge analysis, the approach of the top team in sub-challenge 1 (smallest test set Root Mean Squared Error) was applied to predict time to delivery. **Sub-challenge 2**: Participating teams submitted risk prediction algorithms designed to use as input omics data at two or more time points from individual women coupled with outcome information (control, sPTD or PPROM) for a subset of them (training set), and return disease risk scores for women with blinded outcomes (test set). The algorithms were applied to 70 training/test pairs of datasets (see **Table 1**) to assess within- and across-cohort predictions of preterm birth by whole blood transcriptomics and within-cohort prediction by multi-omics data. Predictions were assessed by Area Under the Receiver Operating Characteristic and Area Under the Precision-Recall curves and aggregated across datasets and prediction scenarios (see **Methods**).

## Results

### Prediction of gestational age by maternal whole blood transcriptomics

We have generated and shared with the community exon-level gene expression data profiled in 703 maternal whole blood samples collected from 133 women enrolled in a longitudinal study at the Center for Advanced Obstetrical Care and Research of the Perinatology Research Branch, NICHD/NIH/DHHS; the Detroit Medical Center; and the Wayne State University School of Medicine. The patient population included women with a normal pregnancy who delivered at term (≥37 weeks) (Controls, N=49), women who delivered before 37 completed weeks of gestation by spontaneous preterm delivery with intact (sPTD, N=34) or ruptured (PPROM, N=37) membranes, and women who experienced an indicated delivery before 34 weeks due to early preeclampsia (N=13) (Dataset Detroit_HTA **Fig. 2a**). After including data from 16 additional normal pregnancies from the same population^23^ (Dataset GSE1139, 32 transcriptomes), the resulting set of 149 pregnancies (see demographic characteristics in **Table S1**), totaling 735 transcriptomes, was divided randomly into training (n=367) and test (n=368) sets; the latter set excludes publicly available data to avoid the possibility that the models are trained with data to be used for testing (**Fig. S1**). The research community was challenged to use data from the training set to develop gene expression prediction models for gestational age, as defined by the last menstrual period (LMP) and ultrasound fetal biometry, and to make predictions based only on gene expressions in the test set. The clinical diagnosis and sample-to-patient assignments were not disclosed to the challenge participants, while gestational age at the time of sampling was also blinded for the test set. Teams were allowed to submit up to 5 predictions for the test samples, and the best submission (smallest Root Mean Squared Error, RMSE) was retained for each unique team. We received 331 submissions for this sub-challenge from 87 participating teams, of which 37 teams provided the required details on the computational methods used to be qualified for the final team ranking in this sub-challenge (**Table S2**).

**Figure 2.**
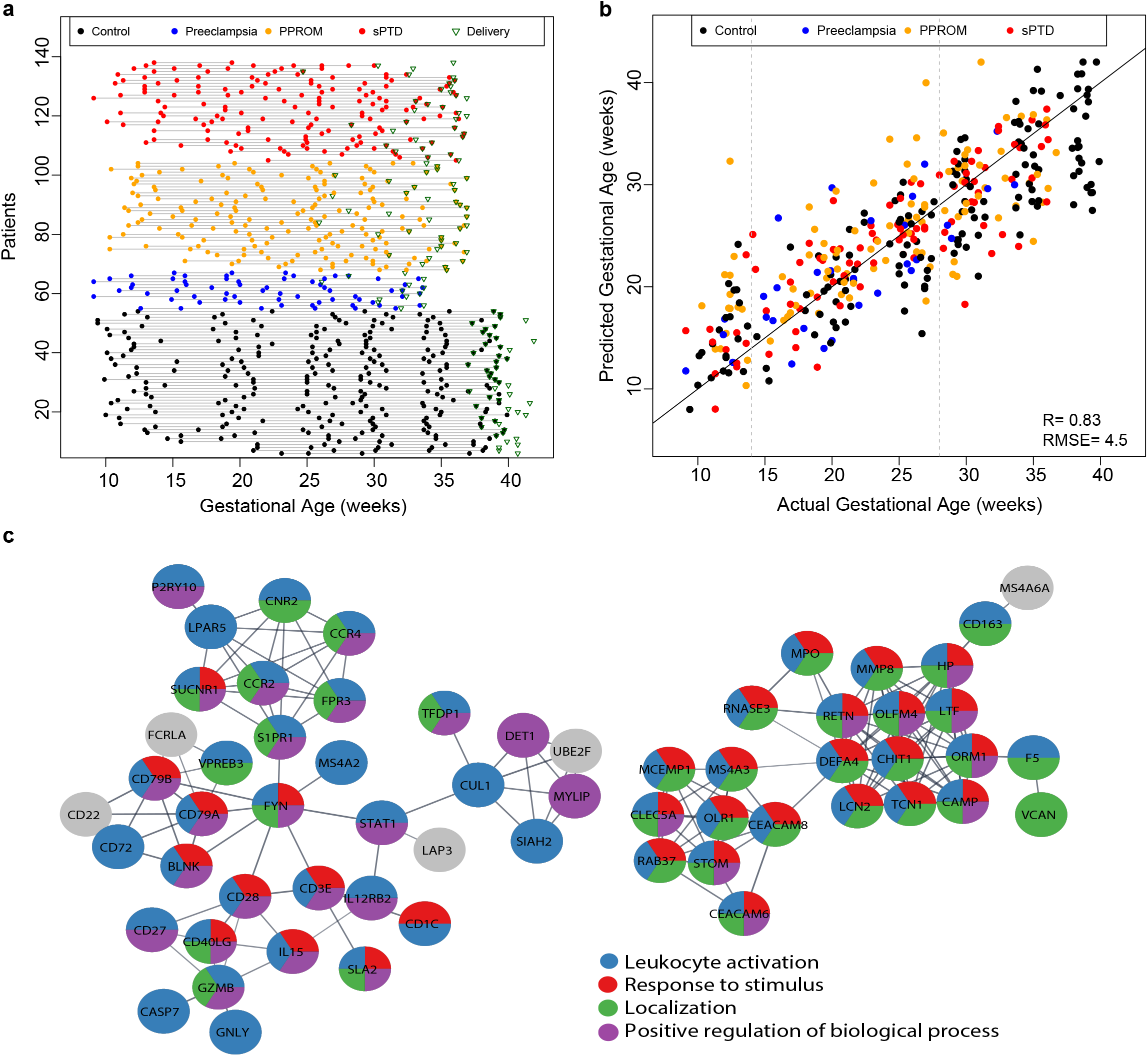
Prediction of gestational age by whole blood transcriptomics. A) Detroit cohort whole blood transcriptomics study design. Each line corresponds to one patient and each dot represents a sample. Gestational ages at delivery are marked by a triangle. B) Test set prediction of gestational age by the model of the top-ranked team (M_GA_Team1). Samples are colored according to the phenotypic group of patients. R: Pearson correlation coefficient; RMSE: Root Mean Squared Error. C) Protein-protein interaction network modules for genes part of the 249-gene core transcriptome predicting gestational age (M_GA_Core). A select group of biological processes enriched among these genes are shown in the pie charts.

Team ranking robustness analysis (see **Methods**) suggested that the predictions of the first-ranked team were significantly better (Bayes factor > 3) than those of the second- and third-ranked teams (**Fig. S2**). Among the top 20 teams, the most frequent methods used to select predictor genes included univariate gene ranking and meta-gene building via principal components analysis as well as literature-based gene selection. Common prediction models included neural networks, random forest, and regularized regression (LASSO and ridge regression), with the latter being used by the top ranked team in this sub-challenge.

The model generated by the first-ranked team in sub-challenge 1 (Team 1) predicted the test set’s gestational ages at blood draw with an RMSE of 4.5 weeks (Pearson correlation between actual and predicted values, r=0.83, p<0.001) (**Fig. 2b**). This prediction model (M_GA_Team1) was based on ridge regression, and the predictors were meta-genes derived by using principal components from expression data of 6,106 genes. As shown in **Fig. 2b**, the gestational age predictions showed little bias in second trimester (14-28 weeks) samples (mean error 0.6 weeks); however, gestational ages predicted of first trimester samples were overestimated (mean error 3.7 weeks) while the third-trimester samples were underestimated (mean error −1.96 weeks). This finding can be understood, in part, by the larger number of second trimester samples relative to first- and third trimester samples, available for training of the model. Of interest, the prediction errors for complicated pregnancies were similar to those of normal pregnancies (ANOVA, p>0.1), suggesting that this model, in general, was robust to obstetrical disease- and parturition-related perturbations in gene expression data (**Fig. S3**).

To identify a core transcriptome predicting gestational age in normal and complicated pregnancies that captures most of the predictive power of the full model (M_GA_Team1) that involved >6000 predictor genes, we combined linear mixed effects modeling for longitudinal data to prioritize gene expressions and then used these as input in a LASSO regression model. The resulting 249 gene regression model (M_GA_Core) (**Fig. S4, Table S3**) had an RMSE=5.1 weeks (Pearson correlation between actual and predicted values, r=0.80) and involved two tightly connected modules related to immune response, leukocyte activation, inflammation- and development-related Gene Ontology biological processes (**Fig. 2c**, **Table S4**). We previously reported that several member genes of these networks (e.g. MMP8, CECAM8, and DEFA4) were most highly modulated in the normal pregnancy group used herein^29^, and others have shown the same to be true at a cell-free RNA level in a Danish cohort^17^. In addition, these data are consistent with the concept that pregnancy is characterized by a systemic cellular inflammatory response^30-34^. In this study, we also show that these mediators correlate with gestational age in both normal and complicated pregnancies, and the latter group contributed more than half of the transcriptomes used to fit and evaluate the models (**Table S1**).

### Comparison of gene expression models and clinical standard in predicting time to delivery in women with spontaneous term or preterm birth

To enable a direct comparison with a previous landmark study of pregnancy dating by targeted cell-free RNA profiling^17^, we used the same methods as described above for model M_GA_Team1, except for the use of a time variable defined backward from delivery and, hence, independent of LMP and ultrasound estimations [Time to delivery, TTD=Date at sample – Date at delivery (weeks)], as response. As in the study by Ngo et al^17^, only those patients with spontaneous term delivery were included in this analysis, thus omitting even the subset of normal pregnancies that had been truncated by elective cesarean delivery. The prediction performance on the test set of the resulting model (M_sTD_TTD) was compared to the LMP and ultrasound fetal biometry, with the latter predicting delivery at 40 weeks of gestation. As shown in **Fig. 3a**, the gene expression model significantly predicted time to delivery with the same accuracy (RMSE, 4.5 weeks, r=0.86, p<0.001) as when predicting LMP and ultrasound-based gestational age in the full cohort of normal and complicated pregnancies. The test set accuracy of the gene expression model, defined as predicting delivery within one week of the actual date^35^, was 45% (5/11) based on third trimester samples, which is comparable to the LMP and ultrasound estimate based on first or second trimester fetal biometry (55%). Of note, 45% accuracy was also reported by Ngo et al^17^ using cell free RNA based on second and third trimester samples.

**Figure 3.**
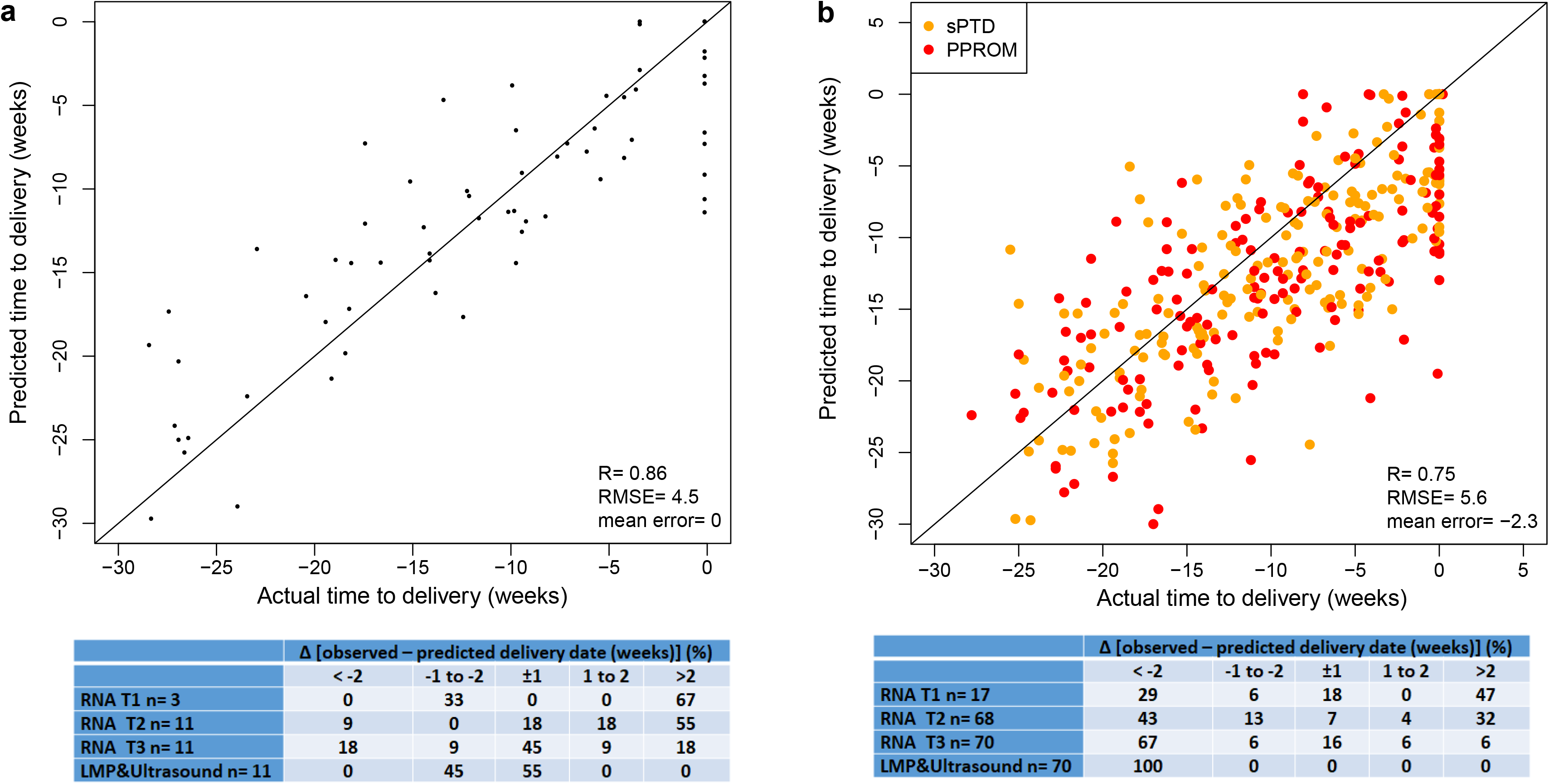
Prediction of time to delivery by whole blood transcriptomics. A) The top panel shows the test set time-to-delivery (TTD) estimates from the M_sTD_TTD model plotted against actual values. The bottom panel shows the distribution of prediction errors (TTD observed - TTD predicted). A negative error means that delivery occurred sooner than expected/predicted, while positive values indicate the opposite. TTD was estimated using RNA measurements from the first- (T1), second- (T2), and third- (T3) trimester samples separately. For comparison, trimesters are defined as in Ngo et al^12^. T1: <12 weeks; T2 = 12-24 weeks, and T3 = 24-37 weeks of gestation. B) Prediction of time-to-delivery in women with spontaneous preterm birth by a gene expression model established in women with spontaneous term delivery (M_sTD_TTD).

When data from all pregnancies with spontaneous term delivery were used to train a transcriptomic model of time to delivery, and apply it to data from women with spontaneous preterm birth, the prediction was found to be statistically significant. However, the error increased (RMSE=5.6) (**Fig. 3b**) relative to the estimate (RMSE=4.5) for prediction of time to delivery in women with spontaneous term delivery (**Fig. 3a**). The additional preterm parturition-specific perturbations in gene expression explain, in part, the added uncertainty in prediction estimates of TTD in spontaneous preterm birth cases compared to spontaneous term pregnancies. Moreover, as expected, the term pregnancy TTD model overestimated the duration of pregnancy of women who were destined to experience preterm birth (**Fig. 3b**). The overestimation (mean prediction error) was 2.3 weeks compared to the five-week gap between the LMP and ultrasound-based gestational ages at delivery in the term (mean, 39 weeks) and preterm (mean, 34 weeks) birth groups. Based on data of 1 to 4 samples collected at 24-37 weeks of gestation for each woman with preterm birth, the M_sTD_TTD model identified 33% of pregnancies to be at risk of delivery within 2 weeks or already past due, with one-half of these, 16% (11/70), predicted to deliver ±1 week from the actual date. This finding contrasts with the LMP-and-ultrasound method, which predicted no cases delivering within ±1 or ±2 weeks of the actual delivery date. This evidence further suggests that the M_sTD_TTD model captured both gene expression changes related to immune- and development-related processes establishing the age of pregnancy as well as the effects of the common pathway of parturition^36,37^. Hence, it generalized to the set of women with spontaneous preterm birth when samples at or near delivery were included and genome-wide gene expression data were available.

### Prediction of preterm birth by maternal blood omics data collected in asymptomatic women (sub-challenge 2)

In this study, we demonstrated that a transcriptomic model of maternal blood derived from data of women with spontaneous term delivery (M_sTD_TTD) can predict spontaneous preterm birth when the data has been collected early in pregnancy as well as near or at the time of preterm parturition up to less than 37 weeks. With sub-challenge 2 of the DREAM Preterm Birth Prediction Challenge, we addressed the more difficult task of predicting preterm birth from data collected up to 33 weeks of gestation while the women were asymptomatic. Of importance, the development of interventions to prevent preterm birth requires pregnant women at risk to be identified as early as possible before the onset of preterm parturition. Moreover, to enable future targeted studies of candidate biomarkers, the maximum number of predictor genes that participating teams could use as predictors in this sub-challenge was limited to a total of 100 genes for prediction of both phenotypes of preterm birth: sPTD and PPROM.

After the first phase of sub-challenge 2 in which predictions were optimistically biased, given the confounding effects of the gestational ages at sampling (see **Methods**), we drew from the Detroit cohort longitudinal study (**Fig. 4a**) only samples collected at specific gestational-age intervals while women were asymptomatic (i.e., prior to an eventual diagnosis of spontaneous preterm delivery or PPROM). Two scenarios of prediction of preterm birth were devised to include cases and controls with available samples collected at the 17-23 and 27-33 weeks (**Fig. 4a**); a third scenario involved patients with available samples collected at three gestational age intervals (17-22, 22-27, and 27-33 weeks) (**Fig. 4b**). The selection of the 17-23 and 27-33 week intervals enabled cross-study model development and testing with the microarray gene expression study of Heng et al.^28^ derived from a cohort in Calgary, Canada. Furthermore, we also included the profiles of 1,125 maternal plasma proteins measured by using an aptamer-based technique^38,39^ in samples collected at 17-23 and 27-33 weeks of gestation from 66 women prior to the diagnosis of preterm birth (62 sPTD and 4 PPROM). These samples were profiled in the same experimental batch with samples from 39 normal pregnancies that we previously described^22,40^, which herein served as controls (**Fig. 4c**). The characteristics of pregnancies with available proteomics profiles are shown in **Table S1**.

**Figure 4.**
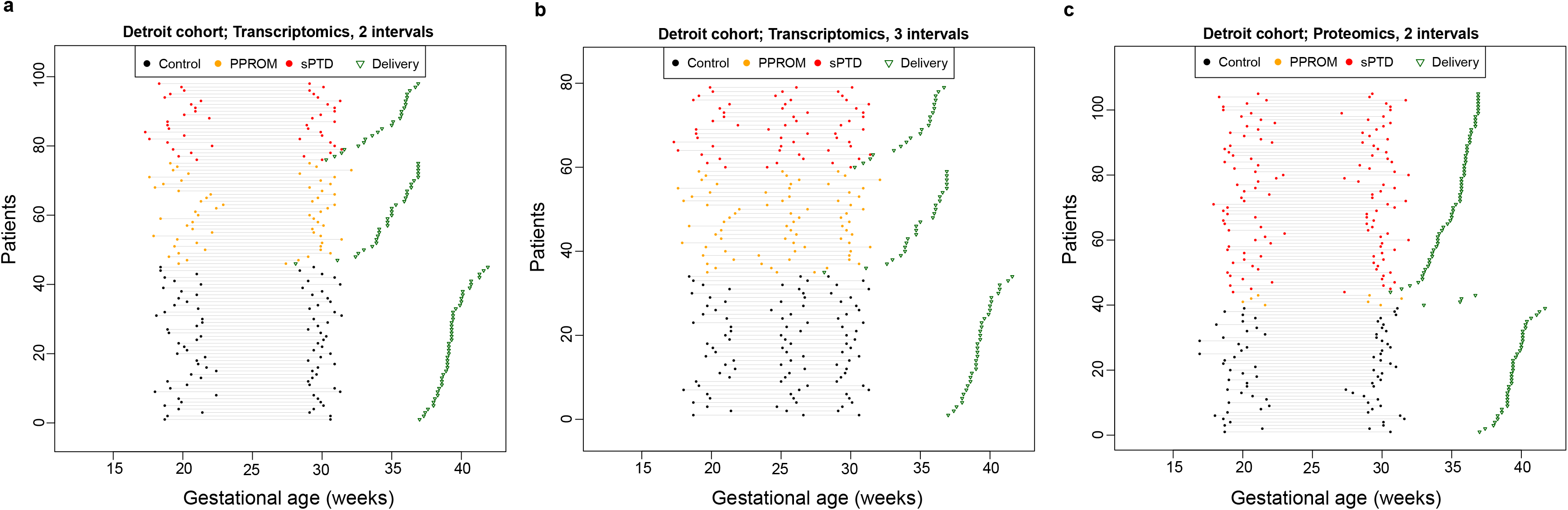
Prediction of preterm birth in asymptomatic women by Detroit omics datasets. From the transcriptomic study in **Fig. 2A**, only controls and preterm birth groups were included. A) Cases were selected if they were asymptomatic at 17-23-week and 27-33–week intervals or B) at 17-22-week, 22-27-week, and 27-33-week intervals. C) Plasma proteomic samples in cases and controls at 17-23-week and 27-33-week intervals.

The prediction algorithms generated by 13 teams who participated in the second phase of sub-challenge 2 were applied by the Challenge organizers to train and test models on 70 pairs of training/test datasets generated under 7 scenarios (**Table 1**). The scenarios differed in terms of omics data type, number of longitudinal measurements per patient, the outcome being predicted, and the patient cohorts used for training/testing (**Table 1**). In all cases, there were no differences in terms of number of samples and gestational age at sampling between the cases and controls (**Fig. 4**).

**Table 1:** Scenarios of spontaneous preterm birth model training and testing using multi-omics data. *Subjects in the original cohort were randomly split into equally sized groups that were balanced with respect to the phenotypes. ** One-fifth of patients from the Detroit cohort (balanced with respect to the phenotypes) were randomly selected in the training set while the remaining four-fifths were used as the test set. ^+^Training set subjects were sampled with replacement from the original cohort to create different versions of the training set, and the trained model was then applied to the original test cohort. sPTB, spontaneous preterm birth; sPTD, spontaneous preterm delivery with intact membranes; PPROM, preterm prelabor rupture of membranes.

| Scenario | Platform | Sample GA (weeks) | Train set | Test set | Number of datasets | Outcomes predicted |
| --- | --- | --- | --- | --- | --- | --- |
| C2 | Transcriptomics | 17-23 & 27-33 | Calgary | Calgary | 10* | sPTD, PPROM |
| D2 | Transcriptomics | 17-23 & 27-33 | Detroit | Detroit | 10* | sPTD, PPROM |
| D3 | Transcriptomics | 17-22 & 22-27 & 27-33 | Detroit | Detroit | 10* | sPTD, PPROM |
| CtoD | Transcriptomics | 17-23 & 27-33 | Calgary | Detroit | 10<sup>+</sup> | sPTD, PPROM |
| DtoC | Transcriptomics | 17-23 & 27-33 | Detroit | Calgary | 10<sup>+</sup> | sPTD, PPROM |
| CDtoD | Transcriptomics | 17-23 & 27-33 | Calgary + Detroit | Detroit | 10** | sPTD, PPROM |
| DP2 | Proteomics | 17-23 & 27-33 | Detroit | Detroit | 10* | sPTD, sPTB |

To assess the prediction performance of the test set in sub-challenge 2, we used both the Area Under the Receiver Operating Characteristic Curve (AUROC) as well as the Area Under the Precision-Recall Curve (AUPRC), the latter being especially suited for imbalanced datasets, e.g., the proteomics set that features more cases than controls (**Fig. 4c**).

**Figure 5** depicts the resulting 28 prediction performance scores (7 scenarios x 2 outcomes x 2 metrics) for each team after the conversion of AUROC and AUPRC metrics into Z scores. Final team rankings were obtained by aggregating the ranks over all prediction performance scores that were significant, according to at least one team, after multiple testing correction (**Table S5**). A rank robustness analysis (**Fig. S5**) determined that the first-ranked team outperformed the second-ranked team and that the second- and third-ranked teams outperformed the fifth-ranked team (Bayes factor > 3). For all scenarios (**Table 1**), the models of Team 1 involved data from 50 genes collected at the last available measurement (closest to delivery), while Team 2 used data collected at the last two available time points for 50 genes selected based on overall expression as opposed to correlation with the outcome. Among other differences in their approaches, Team 1 treated the outcome as a binary variable while Team 2 used a continuous variable derived from gestational age at delivery (see **Methods**). Of note, the top two ranked teams were the same in both sub-challenges 1 and 2.

**Figure 5.**
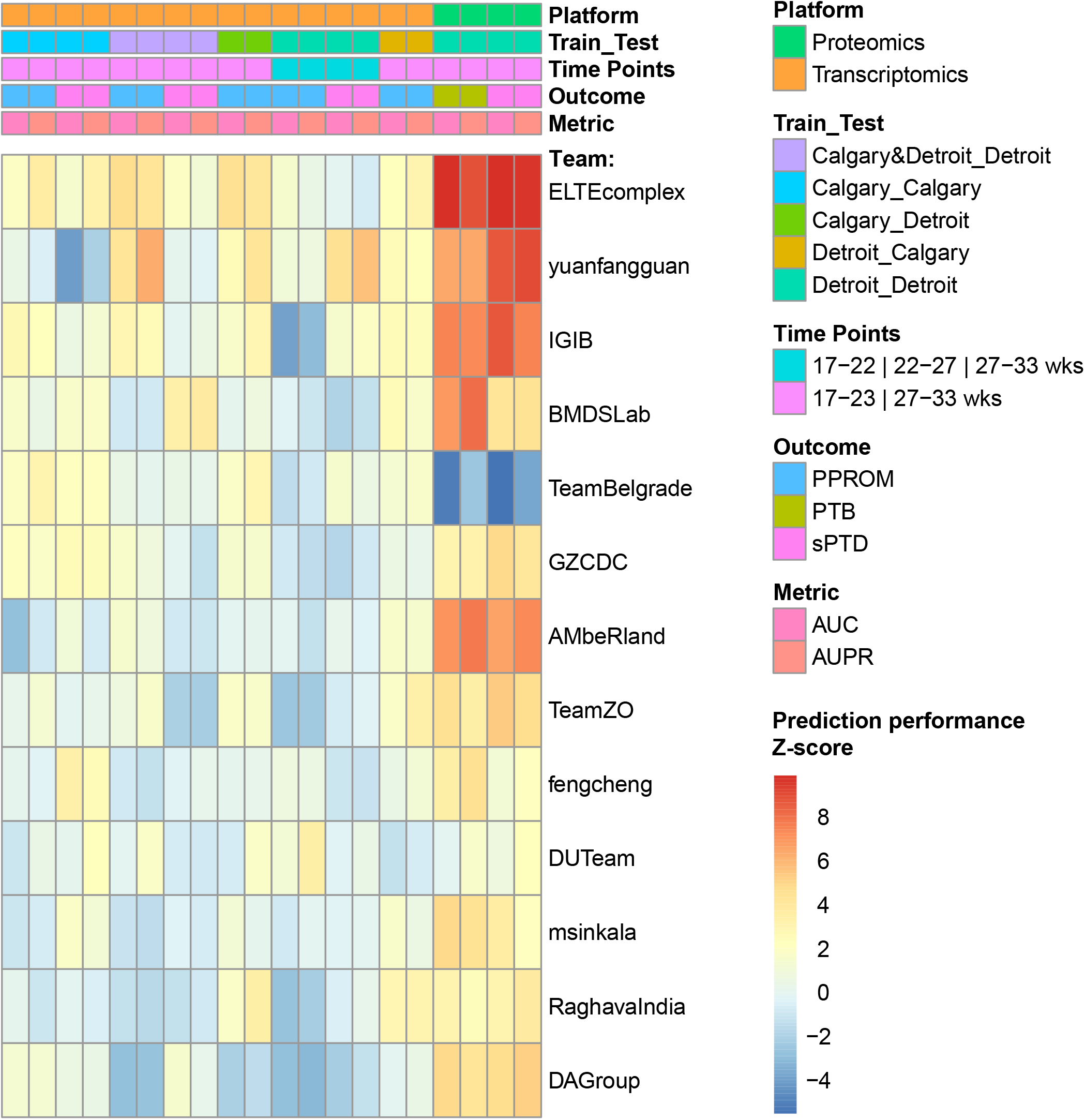
Prediction performance and team ranking in sub-challenge 2. Prediction performance for preterm birth of the approaches of 13 teams under 7 scenarios (Table 1). AUROC and AUPRC metrics were converted into Z-scores and shown as a heatmap.

As depicted in **Fig. 6a**, with the approach of Team 1, one transcriptomic profile at 27-33 weeks of gestation from asymptomatic women predicted PPROM across the cohorts and microarray platforms [AUROC=0.6 (0.56-0.64)] when training the model on the Calgary cohort and testing on the Detroit cohort. Nearly the same performance [AUROC=0.61 (0.56-0.66)] was observed when one-fifth of the Detroit set was added to the Calgary cohort so that microarray platform and cohort effects can be accounted for during the data preprocessing and model training. Prediction of PPROM when training on the Detroit cohort and testing on the Calgary cohort was shy of significance [AUROC=0.54 (0.5-0.57)] based on one sample collected at the 27-33 weeks interval but increased to an AUROC of 0.6 (0.56-0.63); if, in a post-challenge analysis, borrowing from the approach of Team 2, for the same predictor genes, the data from both time points (17-22 and 27-33 weeks of gestation) were used as independent predictors in the model of Team 1 (**Fig S6**). Although separate differential expression analyses of the data from each cohort and time point failed to reach statistical significance after multiple testing correction, the consistency across cohorts and time points of gene expression changes preceding the diagnosis of PPROM was demonstrated by an individual patient data meta-analysis, which identified 402 differentially expressed genes after adjusting for cohort and time point (moderated t-test q-values <0.1) (**Fig. 6b**, **Table S6**). A highly connected protein-protein interaction sub-network corresponding to genes significant in this meta-analysis is shown in **Fig. 6c**, illustrating some of the Gene Ontology biological processes significantly enriched in PPROM and included vesicle-mediated transport and leukocyte-(myeloid and lymphocyte) mediated immunity, among others (**Table S7)**. These data are consistent with the hypothesis that circulating myeloid (monocytes and neutrophils) and lymphoid (T cells) cells are especially activated in women who experience pregnancy complications such as preterm labor^41–44^ and PPROM^45^.

**Figure 6.**
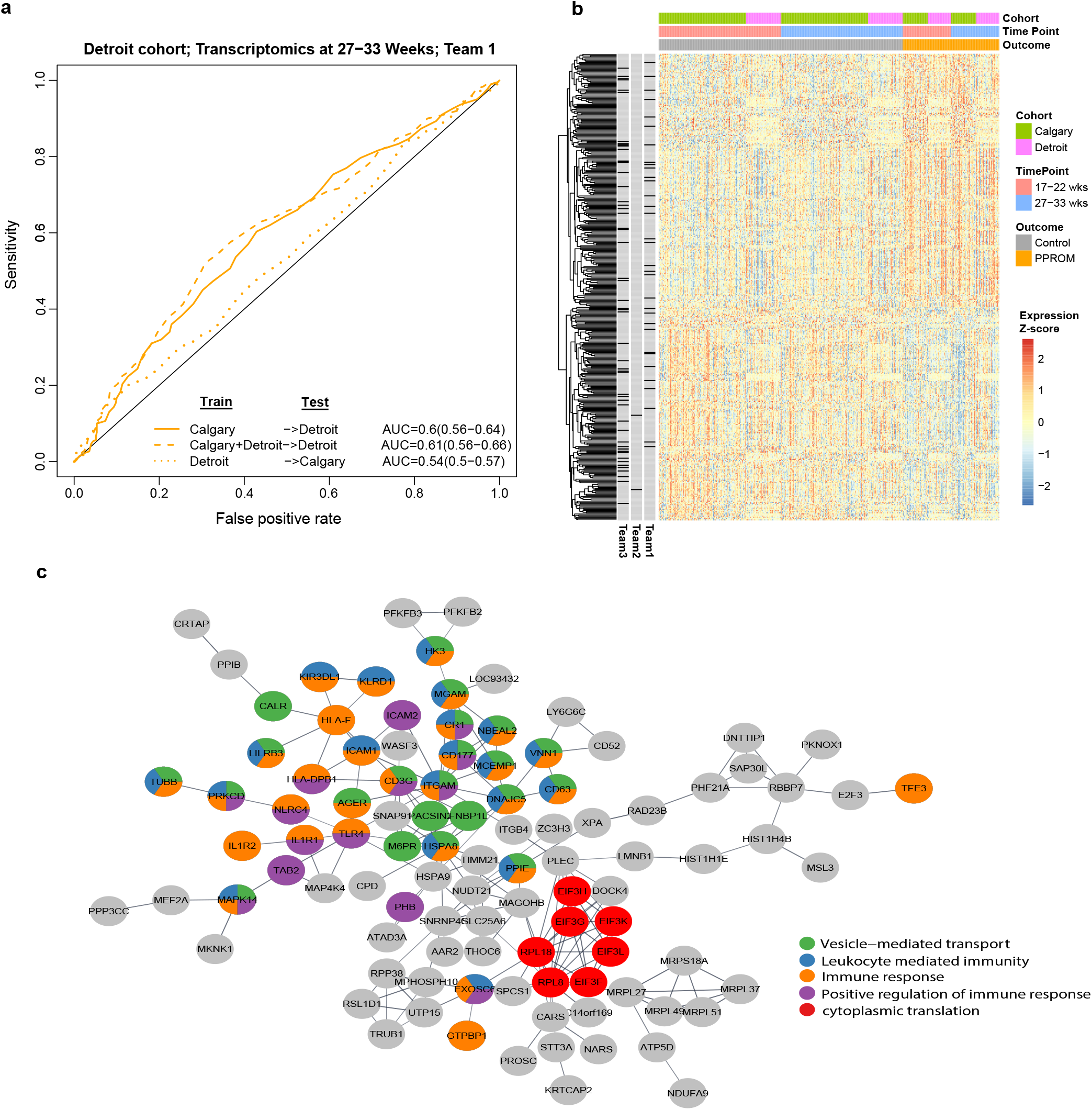
Prediction of preterm prelabor rupture of the membranes from samples collected in asymptomatic women. A) Receiver operating characteristic curve (ROC) representing prediction of preterm prelabor rupture of the membranes (PPROM) by 50 genes across the cohorts and microarray platforms using the Team 1 approach. B) Heatmap of 402 genes differentially expressed in PPROM across the cohorts and time points. Bars on the left indicate gene inclusion as a predictor by the methods of the top 3 teams in sub-challenge 2. C) STRING network constructed from among the 402 genes with differential expression in PPROM. Significantly enriched biological processes are highlighted.

Unlike PPROM, changes in the maternal whole blood transcriptome preceding spontaneous preterm delivery with intact membranes were not shared across cohorts at either the 17-23 or 27-33 week intervals. Within the Detroit cohort, prediction of spontaneous preterm delivery was best performed by Team 2, whose approach leveraged information from the last-available two time points; the prediction performance was significant [AUROC=0.66 (0.6-0.73)] if the data collected at 22-27 and 27-33 weeks of gestation were used, and it was not significant if the data collected at 17-22 and 27-33 weeks of gestation were used instead (**Fig. S7**). This finding is in agreement with previous observations that that closer the sampling to the clinical diagnosis, the higher the predictive value of the biomarkers^40,46,47^. Of interest, 20 of the 50 most highly expressed genes in the Detroit cohort, chosen as predictors by Team 2, showed a significant correlation with gestational age at delivery at both time points simultaneously, suggesting a within-cohort-across-time-point consistency in gene expression changes with gestational age at delivery (q<0.1) (**Table S8**).

Although participating teams in this sub-challenge did not have access to the longitudinal plasma proteomics data in preterm birth included herein when they developed prediction algorithms, their algorithms, when applied to the plasma proteomics set (**Fig. 4c**), resulted in models with test prediction performances that surpassed those obtained using transcriptomic data (**Fig. 5** and **Table S5** and **Fig. 7a**). Prediction of spontaneous preterm delivery by the approach of Team 1 involved 50 plasma proteins selected by random forest model importance from the panel of 1,125 available proteins. The test set accuracy was the highest when using data collected at 27-33 weeks of gestation [AUROC=0.76 (0.72-0.8)] (**Fig. 7A, Table S9**). However, importantly, even one proteome profile at 17-22 weeks of gestation predicted significantly spontaneous preterm delivery [AUROC=0.62 (0.58-0.67)] (**Table S10**), suggesting that this approach has value in the early identification of women at risk. The addition of four cases with PPROM to those with spontaneous preterm delivery did not impact the prediction performance of the proteomics models of Team 1, suggesting that this approach could generalize to both preterm birth phenotypes. Indeed, the increase in plasma protein abundance of PDE11A and ITGA2B preceded the diagnosis of both spontaneous preterm delivery and PPROM in the Detroit cohort at 27-33 weeks of gestation (**Fig. 7b,c**). The tightly interconnected network of proteins built from among those with differential profiles with spontaneous preterm delivery in asymptomatic women at 27-33 weeks of gestation (q<0.1, **Fig. 7b** and **Table S9**) included not only several previously known markers of preterm delivery (IL6, ANGPT1) but also MMP7 and ITGA2B, which we previously described as dysregulated in women with preeclampsia^46^. Member proteins of this network perturbed prior to a diagnosis of spontaneous preterm delivery are annotated to biological processes such as regulation of cell adhesion, response to stimulus, and development (**Fig. 7d**).

**Figure 7.**
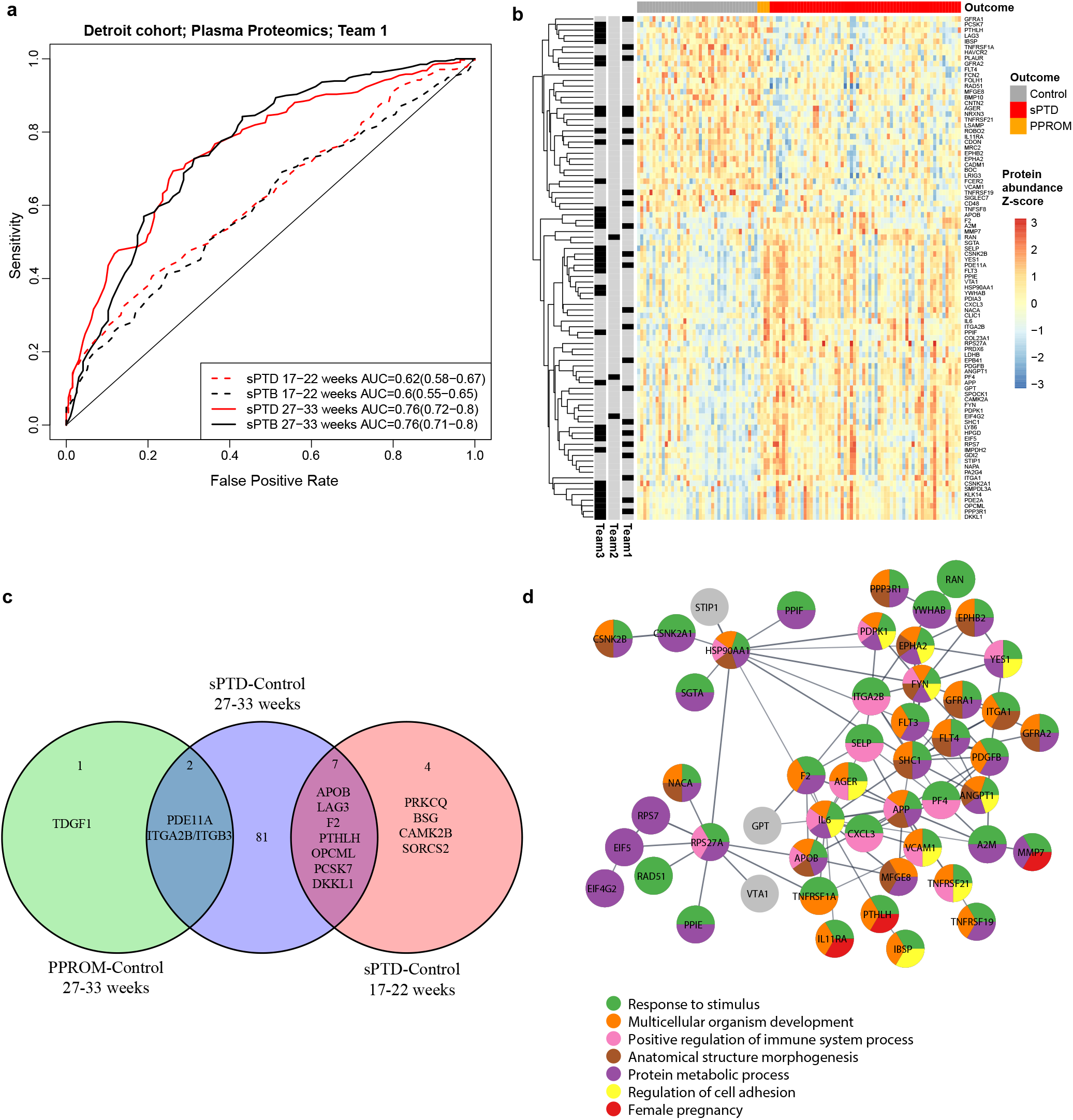
Prediction of spontaneous preterm delivery by plasma proteomic data. A) Receiver operating characteristic (ROC) curve represents the predictions of spontaneous preterm delivery (sPTD) and spontaneous preterm birth (sPTB) [which includes sPTD and preterm prelabor rupture of the membranes (PPROM)] for Team 1. B) Plasma protein abundance for all proteins deemed as significant according to a moderated t-test (q-value<0.1), of which those selected by the top teams as predictors in their models are marked on the left side of the heatmap. C) Overlap of protein changes in those with sPTD at an earlier time point (17-22 weeks of gestation) and with PPROM at 27-33 weeks of gestation. D) Network of proteins among those shown in panel B: each protein node is annotated to biological processes based on corresponding gene ontology.

Given that differences in the patient characteristics could have contributed to the higher prediction performance of spontaneous preterm delivery by plasma proteomics as compared to maternal whole blood transcriptomics, the approach of Team 1 was also evaluated via leave-one-out cross validation on a subset of 13 controls and 17 spontaneous preterm delivery cases for which both types of data originated in the same blood draw. The prediction performance for spontaneous preterm delivery by plasma proteomics remained high [AUROC=0.86 (0.7-1.0)], while prediction by transcriptomic data remained non-significant (**Fig. 8**), hence confirming the superior value of proteomics relative to transcriptomics for this endpoint. Of note, for a fixed number of 50 predictors allowed, a stacked generalization^48^ approach combining predictions from individual platform models via a LASSO logistic regression led to higher leave-one-out cross-validation performance estimate [AUROC=0.89 (0.78-1.0)] compared to building a single model from the combined transcriptomic and proteomic features (**Fig. 8**).

**Figure 8.**
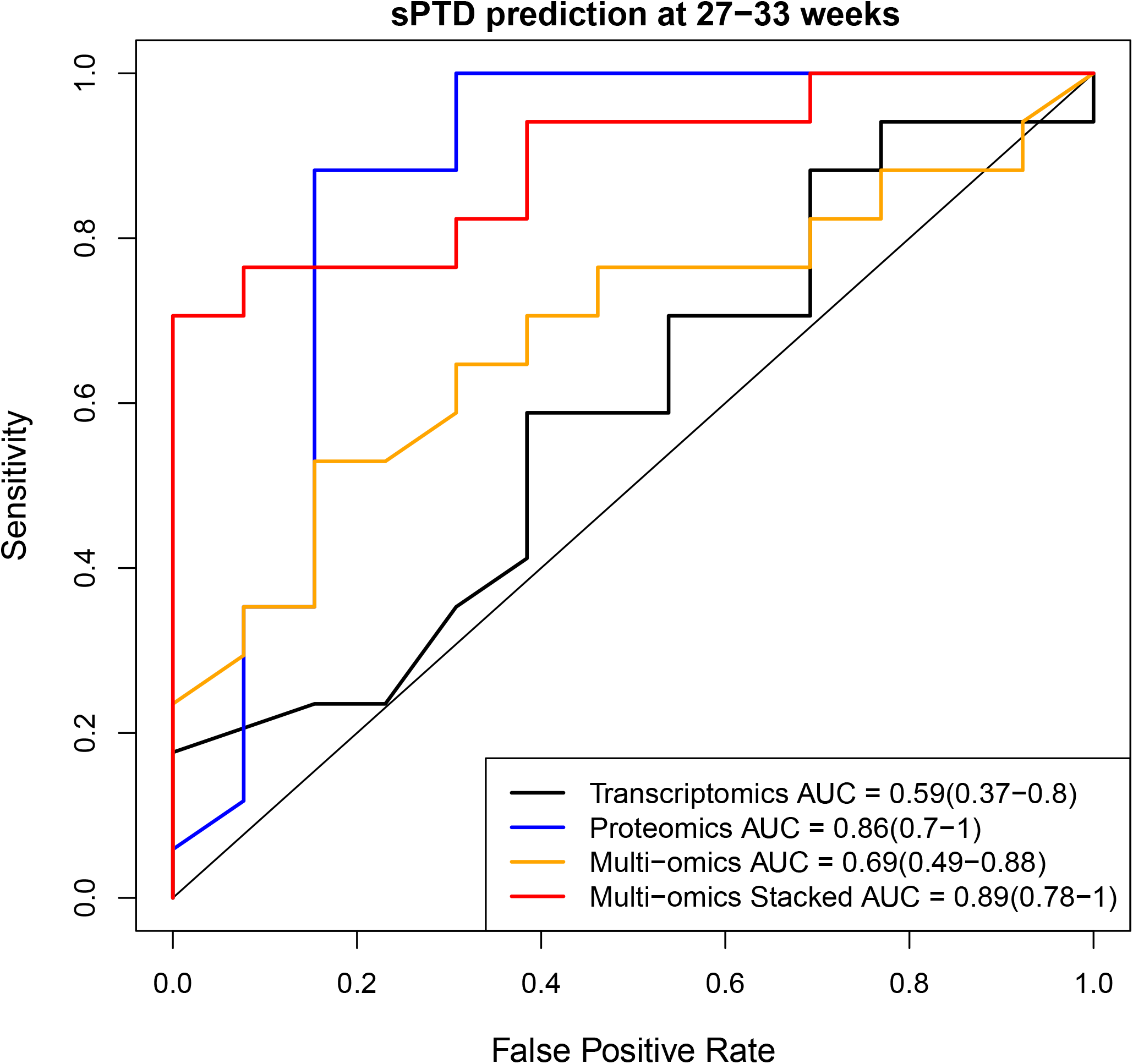
Comparison of prediction performance of spontaneous preterm delivery between platforms. Receiver operating characteristic curve (ROC) for prediction of spontaneous preterm delivery by models obtained with the approach of Team 1 based on a subset of samples for which data from both platforms were available. The multi-omics model was obtained applying the same approach on a concatenated set of proteomic and transcriptomic features. The multi-omics stacked generalization approach involved combining predictions from models based on each platform via logistic regression.

To extract further insight into the computational approaches best suited to predict preterm birth from longitudinal omics data in sub-challenge 2, we investigated which computational aspects explained the higher performances of the top two teams. Given that Team 1 relied only on omics data at the last available time point (T2), we kept all aspects of its method except for the temporal information considered among the following: a) first point (T1), b) change in expression between T2 and T1 (slope), or c) a combined approach in which slopes for all genes and measurements at T2 compete for inclusion in the 50 allowed predictors for a given outcome (PPROM or spontaneous preterm delivery). As shown in **Fig. S8,** none of these approaches would have improved prediction performance relative to the baseline approach that only considered data from the last time point (T2). We then considered several key aspects of the approach of Team 2 and have subsequently incorporated them in the approach of Team 1 to determine whether such hybrid approaches could translate into higher performances relative to the baseline approach. In particular, we have modified the approach of Team 1: a) to start with only the top half of the most highly abundant features on each platform, b) to convert the binary classification (preterm versus term) into a regression of gestational age at delivery, and c) given the selected 50 predictor genes based on the correlation of T2 expression values with the outcome, to add the expression of those gene at the previous time point as independent predictors in the random forest model. Of these three scenarios, the last, which expands the number of predictors from 50 to 100 without increasing the number of unique genes, slightly outperformed the approach of Team 1’s overall prediction scenarios (**Fig S8**) and led to the consistent prediction of PPROM in all cross-study analyses (see improvement in prediction from **Fig. 6a** to **Fig. S6**). Interestingly, simply doubling the number of genes measured at T2 that were allowed as predictors in the model (from 50 to 100) led to a worse overall prediction performance relative to the approach of Team 1 that used only 50 genes at T2 (**Fig. S8**). This finding suggests that for preterm birth prediction, it is more important to measure the right markers at one additional time point than to double the number of markers at the most recent time point.

## Discussion

In this study, we have evaluated maternal blood omics data to predict gestational age and the risk of preterm birth. Although the main interest herein was the prediction of spontaneous preterm birth, the correlation of omics data with advancing gestation was relevant not only to serve as a positive control for the evaluation of omics data, but also to possibly provide relevant information for the development of more affordable tools to date pregnancy. We chose the DREAM collaborative competition framework^27^ to identify the best computational methods for making inferences and to assess them in an unbiased and robust way based on longitudinal omics data that we and others have generated. DREAM Challenges have been used to establish unbiased performance benchmarks across a wide array of prediction tasks^49-53^. Moreover, the results gained from these Challenges define community standards and advancements in many scientific fields^54,55^.

Collectively, sub-challenge 1 and the additional post-challenge analyses demonstrated that models based on the maternal whole blood transcriptome i) significantly predict LMP and ultrasound-defined gestational age at venipuncture in both normal and complicated pregnancies (RMSE=4.5), ii) predict a delivery date within ±1 week in women with spontaneous term delivery with an accuracy (45%) comparable to the clinical standard (55%), and iii) outperform the latter approach for timing of delivery in women who are destined to experience spontaneous preterm birth (33% predicted within ±2 weeks of delivery versus 0% for the LMP and ultrasound method). Of interest, the accuracy of dating gestation in women with spontaneous term delivery was similar to the report by Ngo et al.^17^ who used cell-free RNA profiling in a Danish cohort, although that study involved more frequent (weekly) sampling of fewer genes (about 50 immune-, placental- and fetal liver-specific) instead of the genome-wide data used herein. However, in the study by Ngo et al.^17^, the time-to-delivery transcriptomic model derived from samples of women with normal pregnancy failed to predict delivery dates on independent cohorts of women with preterm birth, while 33% of preterm birth cases herein were accurately identified as being at risk of delivery within ±2 weeks. A possible explanation, in addition to the cohort differences between the training and testing sets in the previous study, is that our model of normal pregnancy captured not only gene changes establishing the gestational age, but also those changes involved in the common pathway of labor. While the prediction of preterm birth by omics data collected up to less than 37 weeks including samples taken when women were symptomatic was demonstrated above without using any data from preterm birth cases to establish the model, it was also previously shown by others who used data from both cases and controls^10,20,56,57^.

In the context of sub-challenge 2 of the DREAM Preterm Birth Prediction Challenge, we have tackled the issue of predicting preterm birth from samples collected while women were asymptomatic prior to 33 weeks of gestation. Overall, although prediction performance was low (AUROC=0.6 at 27-33 weeks of gestation), the sub-challenge and post-challenge analyses provide evidence of changes in maternal whole blood gene expression that precede a diagnosis of PPROM and that are shared across gestational-age time points and cohorts/microarray platforms. However, although some correlations of gene expression to gestational age at delivery were consistent across time points within the Detroit cohort (**Table S8**), the cross-cohort changes preceding spontaneous preterm delivery with intact membranes were not consistent. Moreover, the Detroit within-cohort proteomic results, obtained with methods developed without any tuning to the proteomic data, represent evidence of superior performance by plasma proteomics (AUROC=0.76 at 27-33 weeks) compared to whole blood transcriptomics in the prediction of spontaneous preterm delivery. Although we and other investigators reported the value of aptamer-based SomaLogic assays to predict early^46^ and late preeclampsia^40^, this is the first study conducted to evaluate this platform to predict preterm birth, and we found it to be of superior value as compared to whole blood transcriptomics platform in predicting spontaneous preterm delivery. Two possible limitations to the comparison between platforms are the lower sample size utilized to analyze the same blood draws and the much larger number of transcriptomic than proteomic features, which made the “needle in the haystack” problem more difficult for the transcriptomic platform. This curse of dimensionality was noted when transcriptomic and proteomic features were combined, resulting in a lower performance estimate for the multi-omics model obtained with the approach of Team 1, than for proteomics data alone. Although herein the remedy to this issue was to combine the predictions of each platform into a meta-model (stacked generalization) (**Fig. 8**), alternative approaches focus on biologically plausible sets of features derived by single-cell genomics. This latter category of methods was demonstrated to predict preeclampsia^47,58^ and to distinguish between women with spontaneous preterm labor and the gestational age-matched controls^43,44^.

The use of crowdsourcing to evaluate computational approaches and longitudinal multi-omics data to predict preterm birth is a major strength of this study. The primary advantage of importance is that many perspectives of this problem were implemented by the machine learning community. Coincidently, the first and second best-performing teams were the same for both sub-challenges, which is indicative of the team’s skill as opposed to chance, a fact that has been observed in several other crowd-sourcing initiatives, e.g., sbv IMPROVER^23,59,60^, CAGI^61-64^ and DREAM^50,65,66^. The second advantage of the DREAM Challenge framework is that the model development and the prediction assessment are separate, thus the risk of overstating the prediction performance is reduced. This robust evaluation of prediction performance, combined with a separate consideration of preterm birth phenotypes (spontaneous preterm delivery and PPROM), of time points at sampling, and multi-omic platforms, makes this work one of the most comprehensive longitudinal omics studies in preterm birth. Finally, the work herein has resulted in computational algorithms with implementations made available to the community with an open source license, allowing for reproducible research and applications to other similar research questions based on longitudinal omics data.

## Methods

### Study design

Women who provided blood samples included in the transcriptomic and proteomic studies described in the Results section were enrolled in a prospective longitudinal study at the Center for Advanced Obstetrical Care and Research of the Perinatology Research Branch, NICHD/NIH/DHHS; the Detroit Medical Center; and the Wayne State University School of Medicine. Blood samples were collected at the time of prenatal visits, scheduled at four-week intervals from the first or early second trimester until delivery, during the following gestational-age intervals: 8-<16 weeks, 16-<24 weeks, 24-<28 weeks, 28-<32 weeks, 32-<37 weeks, and >37 weeks. Collection of biological specimens and the ultrasound and clinical data was approved by the Institutional Review Boards of Wayne State University (WSU IRB#110605MP2F) and NICHD (OH97-CH-N067) under the protocol entitled “Biological Markers of Disease in the Prediction of Preterm Delivery, Preeclampsia and Intra-Uterine Growth Restriction: A Longitudinal Study.” Cases and controls were selected retrospectively.

### Clinical definitions

The first ultrasound scan during pregnancy was used to establish gestational age if this estimate was more than 7 days from the LMP-based gestational age. The first ultrasound scan was obtained before 14 weeks of gestation for 70% of the women, and 95% of the women underwent the first ultrasound before 20 weeks of gestation. Preeclampsia was defined as new-onset hypertension that developed after 20 weeks of gestation (systolic or diastolic blood pressure ≥140 mm Hg and/or ≥90 mm Hg, respectively, measured on at least two occasions, 4 hours to 1 week apart) and proteinuria (≥300 mg in a 24-hour urine collection, or two random urine specimens obtained 4 hours to 1 week apart containing ≥1+ by dipstick or one dipstick demonstrating ≥2+ protein)^67^. Early preeclampsia was defined as preeclampsia diagnosed before 34 weeks of gestation, and late preeclampsia was defined by diagnosis at or after 34 weeks of gestation^68^. The diagnosis of PPROM was determined by a sterile speculum examination with documentation of either vaginal pooling or a positive nitrazine or ferning test^69^. Spontaneous preterm labor and delivery was defined as the spontaneous onset of labor with intact membranes and delivery occurring prior to the 37^th^ week of gestation^70^.

### Maternal whole blood transcriptomics

RNA was isolated from PAXgene^®^ Blood RNA collection tubes (BD Biosciences, San Jose, CA; Catalog #762165) and hybridized to GeneChip^™^ Human Transcriptome Arrays (HTA) 2.0 (P/N 902162), as we previously described^29^. Microarray experiments were carried out at the University of Michigan Advanced Genomics Core, a part of the Biomedical Research Core Facilities, Office of Research (Ann Arbor, MI, USA). Raw intensity data (CEL files) were generated from array images using the Affymetrix AGCC software. CEL files from this study and those for the Calgary cohort were preprocessed separately for each platform. ENTREZID gene level expression summaries were obtained with Robust Multi-array Average (RMA)^71^ implemented in the *oligo* package^72^ using suitable chip definition files from http://brainarray.mbni.med.umich.edu. Since samples in the Detroit cohort were profiled in several batches, correction for potential batch effects was performed using the *removeBatchEffect* function of the *limma*^73^ package in *Bioconductor*^74^. Cross-study/platform analyses were performed on a combined dataset after quantile normalizing data across all samples for the set of common genes, followed by platform effect-removal.

### Maternal plasma proteomics

Maternal plasma protein abundance was determined by using the SOMAmer (Slow Off-rate Modified Aptamer) platform and reagents to profile 1,125 proteins^38,39^. Proteomic profiling services were provided by SomaLogic, Inc. (Boulder, CO, USA). The plasma samples were diluted and then incubated with the respective SOMAmer mixes, and after following a suite of steps described elsewhere^38,39^, the signal from the SOMAmer reagents was measured using microarrays. The protein abundance in relative fluorescence units was obtained by scanning the microarrays. A sample-by-sample adjustment in the overall signal within a single plate was performed in three steps per manufacturer’s protocol, as we previously described^22,40^. Outlier values (larger than *2×* the 98^th^ percentile of all samples) were set to 2*×* the 98^th^ percentile of all samples. Data was log2 transformed before applying machine learning and differential abundance analyses.

### Sub-Challenge 1 organization

For sub-challenge 1, aimed at predicting gestational age at sampling from whole blood transcriptomic data in normal and complicated pregnancies, a training set and a test set were generated (**Fig. S1**). Transcriptomic gene expression data were made available to participants for both the training and test sets. Gestational age was provided for the training set and participants were required to submit predicted gestational-age values for the test set, which were compared in real time against the gold standard; the RMSE was posted to a leaderboard that was live from May 22, 2019, to August 15, 2019. Up to five submissions per team were allowed, and they were ranked by the RMSE, and the smallest value was retained as entry for each unique team (**Table S1**). Only the teams who described their approach and provided the analysis code were retained in the final team rankings.

### Sub-Challenge 1 team rank stability analysis

To determine whether differences in gestational-age prediction accuracy between the different teams were substantial, we have simulated the challenge by drawing 1000 bootstrap samples of the test set. RMSE values were calculated for each submission (1 to at most 5) for each team, and we retained the submission with the smallest RMSE. Team ranks were calculated and the Bayes factors were then calculated as the ratio between the number of iterations in which the team *k* performed better than the team ranked next *(k+1)* relative to the number of iterations when the reverse was true. A Bayes factor >3 was considered a significant difference in ranking.

### Sub-Challenge 1 top two algorithms

#### Team 1

The first-ranked team in this sub-challenge (authors B.A.P. and I.C.) used gene-level expression data after filtering out samples considered as outliers, followed by the standardization of gene expression for each microarray experiment batch separately. Genes were ranked by using singular value decomposition, and those genes having higher dot products with singular vectors that correspond to large singular values across the training samples were assigned a higher score. In the next step, ~6000 genes were selected based on the described ranking, which was based on cross-validation results on the training set using a ridge regression model. Ridge regression^75^ models were fitted using the *Sklearn* package in *Python* (version 3).

#### Team 2

The second-ranked team in this sub-challenge (author Y.G.) applied quantile normalization to gene level expression data, followed by the modeling of the gestational-age values using Generalized Process Regression and Support Vector Regression. Model tuning parameters were optimized using a grid search, and predictions by the two approaches were weighted equally. Models were fit using *Octave*.

### Sub-Challenge 2 organization

In the first phase of sub-challenge 2, participants were invited to develop preterm birth prediction algorithms using gene expression data from longitudinal transcriptomic data collected at 17-36 weeks of gestation from women with a normal pregnancy and from cases of preterm birth (spontaneous preterm delivery and PPROM) illustrated in **Fig. 2**. The training set was composed of data from the Calgary cohort and a fraction of the Detroit cohort (**Fig. 2**), while the test set comprised the remainder of the Detroit cohort. Teams were requested to submit a risk value (probability) for all samples when classifying test samples as sPTD versus Control, and as PPROM versus Control. The AUROC and AUPRC were calculated separately for each prediction task and the ranks for each of the resulting four performance measures were calculated for each team and aggregated by summation. Two predictions per team were allowed and performance results on the test set were posted to a live leaderboard from August 15, 2019, to December 5, 2019. Because the prediction models developed in this phase of the Challenge could have captured eventual differences between the cases and controls in terms of the timing and number of samples, a second phase of the Challenge was organized (December 5, 2019 to January 3, 2020) for which teams were asked to provide prediction algorithms (computer code) instead of predictions of a given test dataset. The algorithms were applied as implemented by the participants without any tuning to the 70 pairs of training and test datasets described in **Table 1**. In each of the 7 scenarios in **Table 1**, there were 2 outcomes predicted (sPTD vs Control; and PPROM vs Control), except for proteomic data (scenario DP2), where the feasible comparisons were sPTD versus control and PTB versus control; the PTB group was defined as the union of sPTD and PPROM cases. As in the first phase of the challenge, the AUROC and AUPRC were used to assess predictions for each outcome. The resulting 28 prediction performance scores (7 scenarios x 2 outcomes x 2 metrics) for each team were converted into Z-scores by subtracting the mean and dividing by the standard deviation of these metrics obtained from 1,000 random predictions (random uniform posterior probabilities). Further, only the combinations of scenarios and outcomes resulting in a significant prediction performance (False Discovery Rate-adjusted p-value, q<0.05) for at least one of the 13 teams, were considered further for team ranking, resulting in 20 performance criteria for each team. Teams were ranked by each of the 20 prediction performance criteria (columns in **Fig. 8**), and a final rank was generated based on the sum of the ranks over all criteria (**Table S5**).

### Sub-challenge 2 team rank robustness analysis

To assess the significance of the differences in prediction performance of preterm birth among the teams based on omics data, we have used the same ranking procedure described above in more than 1,000 simulated iterations of the sub-challenge. At each iteration, the rankings were calculated by using prediction performance results that correspond to a bootstrap sample of the 10 train/test instances pertaining to each scenario (**Table 1**) and, at the same time, taking a bootstrap sample of the prediction criteria (columns in the **Fig. 8**). Bayes factors were then calculated as the ratio between the number of iterations in which the team *k* performed better than the team ranked next (*k*+1) relative to the number of iterations when the reverse was true. A Bayes factor >3 was considered a significant difference among rankings.

### Sub-Challenge 2, the top three algorithms

#### Team 1

The algorithm of the first-ranked team in this sub-challenge (authors B.A.P. and I.C.) starts with standardizing the input omics data so that they have a zero mean and a standard deviation of 1 for each omics platform (if more than one in an input set, which was the case while training and testing across the platforms). A random forest classifier with 100 trees was fit to each prediction task (sPTD versus Control and PPROM versus Control). The top 50 features, ranked by importance metric derived from the random forest, were selected for each task separately and used to fit a final model on the training data. Random forest repressors were fitted using the *Sklearn* package in *Python* (version 3).

#### Team 2

The approach of the second-ranked team in this sub-challenge (author Y.G.) first centers the data of each feature around the mean for each platform (if more than one) in a given input set. Then, data is quantile normalized to make identical the distributions of feature data across the samples. Next, the top 50 features with the highest average over all samples are retained, and the feature values for the last-available two time points for each subject are used as predictors (100 predictors) in a Generalized Process Regression model, a Bayesian non-parametric regression technique. The two parameters of GPR regression were preset to an eye value of 0.75, which represents how much noise is assumed in the data, and a sigma of 10, a data normalization factor. Models were fitted using Octave.

#### Team 3

The approach of the third-ranked team in this sub-challenge (author R.K.) starts with the selection of the top 50 features ranked by statistical significance p-value derived from a t-test or Wilcoxon test, depending on the normality of the data, and determined by a Shapiro test. Then, using the selected features, linear, sigmoid and radial Support Vector Machines models are fitted and compared via 5-fold cross validation, and the predictions for the best method were averaged over the five trained models. Models were fit using the *e1071* package^76^ in R.

### Post-challenge differential expression and abundance analyses

Differences in gene expression or protein abundance between the cases and controls were assessed based on linear models implemented in the *limma* package^77^ in Bioconductor. When data across time points and/or cohorts were combined, these factors were included as fixed effects in the linear models. Downstream analyses of the differentially expressed genes involved enrichment analysis via a hypergeometric test implemented in the *GOstats* package^78^ to determine the over-representation of Gene Ontology^79^ biological processes among the significant genes. The background list in the enrichment analyses featured all genes profiled on the microarray platform. For proteomic-based enrichment analyses, protein-to-gene annotations from the manufacturer (SomaLogic) were used as input in the *stringApp* version (1.5.0)^80^ in *Cytoscape* (version 3.7.2)^81^ using the whole genome as background. A false discovery rate adjusted q<0.05 was used in enrichment analyses to infer significance. Networks of high-confidence protein-protein interactions (STRING confidence score > 0.7) were constructed from the lists of significant genes/proteins using *stringApp* in Cytoscape. For visualization, the most interconnected subnetworks were displayed and nodes were annotated to significantly enriched biological processes.

### Identification of a core transcriptome predicting gestational age

To identify a core transcriptome that can predict gestational age in normal and complicated pregnancies, linear mixed-effects models with splines were applied to prioritize genes that change with gestational age while accounting for the possible non-linear relation and for the repeated observations from each individual, as we previously described^29^. Of note, participating teams could have not used such an approach given that sample-to-patient annotations were not provided on the training data. Then, the genes that did not change in average expression by at least 10% over the 10-40-week span were filtered out, and the remaining genes were ranked by p-values from the linear mixed-effects models. The top 300 genes were then used as input in a LASSO regression model (elastic net mixing parameter alpha =0.01) for which the shrinkage coefficient (lambda) was determined by cross-validation, leading to 249 genes with non-zero coefficients in the model (**Table S2**). Of note, using more than 300 genes as input in the ridge regression model did not further reduce the RMSE on the test set. LASSO models were fit using the *glmnet* package^82^ in R.

## Data availability

The transcriptomic and proteomic data from the Detroit cohort described herein is available as a Gene Expression Omnibus super-series (GSE149440) and GSE150167, respectively. They were also submitted to the March of Dimes repository (https://www.immport.org/shared/study/SDY1636).

## Code availability

Analysis scripts for transcriptomic data preprocessing and for building prediction models based on the approaches of the participating teams in sub-challenges 1 and 2 are available from the Challenge website (www.synapse.org/pretermbirth). Direct links to method write-up and computer implementations for prediction of gestational age and preterm birth are also available in **Tables S2** and **Table S5**, respectively. Moreover, R code vignettes demonstrating the use of participant methods and key post-challenge analyses were also provided at www.synapse.org/pretermbirth.

## Acknowledgements

This research was supported, in part, by the Perinatology Research Branch, Division of Obstetrics and Maternal-Fetal Medicine, Division of Intramural Research, *Eunice Kennedy Shriver* National Institute of Child Health and Human Development, National Institutes of Health, U.S. Department of Health and Human Services (NICHD/NIH/DHHS) and, in part, with federal funds from NICHD/NIH/DHHS under Contract No. HHSN275201300006C.

A.L.T. and N.G.-L. were also supported by the Wayne State University Perinatal Initiative in Maternal, Perinatal and Child Health.

R.R. has contributed to this work as part of his official duties as an employee of the United States Federal Government.

The authors acknowledge Alison Paquette for insightful discussions about available maternal omics datasets in preterm birth and Maureen McGerty (Wayne State University) and Corina Ghita for proofreading and copyediting this manuscript.

N.A. was supported by the Bill and Melinda Gates Foundation (OPP1112382 and OPP1113682), the March of Dimes Prematurity Research Center at Stanford University, the Burroughs Wellcome Fund, the National Center for Advancing Translational Sciences, and the Robertson Family Foundation (KL2TR003143).

I.C and B.A.P. were supported by the National Research, Development and Innovation Fund of Hungary (Project No. FIEK_16-1-2016-0005).

Y.G. was supported by grants from the National Institutes of Health (NIH/NIGMS R35GM133346-01) and National Science Foundation (NSF/DBI #1452656).

R.K. was supported by a Senior Research Fellowship Award from the Council of Scientific and Industrial Research of India (HCP00013).

G.A. and M.S. were supported by the March of Dimes.

## Author contributions

A.L.T, R.R., M.S., N.A., G.S., and J.C.C. designed the challenge. A.L.T., T.Y., and J.C.C. developed tools to receive and evaluate participant submissions. The top-performing approach was designed by I.C. and B.A.P. Data analysis for the top performing approach was conducted by B.A.P. The DREAM Consortium provided predictions for sub-challenge 1, as well as method implementations and descriptions for sub-challenge 1 and 2. A.L.T. and B.D. applied method implementations to datasets for sub-challenge 2 and designed and applied hybrid methods between approaches of top two teams. A.L.T., B.D., B.P.A., G.B., G.A. and J.C.C. performed post challenge analyses. A.L.T, R.R., M.S., N.A., J.C.C. and G.S. interpreted the results of the challenge. A.L.T., R.R., N.G.L., T.C., S.S.H. and C.D.H. designed the protocol in which patients were enrolled and coordinated the collection and generation of the clinical and omics data. A.L.T., N.G.L., M.S., J.C.C. wrote the manuscript. A.L.T, R.R., J.C.C. and G.S. supervised the project.

## The DREAM Preterm Birth Prediction Challenge Consortium

Nima Aghaeepour^18^, Gaia Andreoletti^10,11^, Benan Bardak^22^, Madhuchhanda Bhattacharjee^23^, Gaurav Bhatti^1,2^, Michael Blair^24^, Huiyuan Chen^25^, Feng Cheng^26^, Tinnakorn Chiaworapongsa^2^, Changje Cho^27^, Junseok Choe^28^, Mohit Choudhary^29^, James C. Costello^21^, Istvan Csabai^4^, Yang Dai^30^, Ophilia Daniel^31^, Bikram K. Das^32^, Francisco de Abreu e Lima^33^, Anjali Dhall^34^, Bogdan Done^1,2^, Işıksu Ekşioğlu^22^, Bogdan N. Gavrilovic^35^, Nardhy Gomez-Lopez^1,2,14^, Yuanfang Guan^12^, Akshay Gupta^29^, Romeharsh Gupta^29^, Rohan Gurve^29^, Dániel Györffy^36,37^, Sonia S. Hassan^2,16,17^, Eric D. Hill^38^, Chaur-Dong Hsu^1,2,17^, Jinseub Hwang^39^, Yuguang F. Ipsen^40^, Riza Işik^22^, Priyansh Jain^41^, Pratheepa Jeganathan^42^, Sujae Jeong^43^, Chan-Seok Jeong^44^, Anshul Jha^29^, JinZhu Jia^45^, Jaewoo Kang^46^, Hyojin Kang^44^, Gaurang A. Karwande^47^, Harpreet Kaur^48^, Hannah Kim^49^, Keonwoo Kim^28^, Sunkyu Kim^28^, Dohyang Kim^27^, Junseok Kim^27^, Jongtae Kim^39^, Min-Jeong Kim^50^, Amrit Koirala^32^, Adriana N. König^51^, Prachi Kothiyal^52^, Vladimir B. Kovacevic^35^, Aleksandra V. Kovacevic^53^, Shiu Kumar^54^, Chandrani Kumari^55^, Christoph F. Kurz^51,56^, Rintu Kutum^13^, Taeyong Kwon^27^, Thuc D. Le^31^, Kyeongjun Lee^39^, Hyungyu Lee^57^, Dawoon Leem^57^, Shuya Li^58^, Weng Khong Lim^59,60^, Xinyue Liu^61^, Yunan Luo^62^, Bahattin C. Maral^22^, Suyash Mishra^63^, Yeongeun Nam^27^, Leelavati Narlikar^64^, Thin Nguyen^65^, Zoran Obradovic^66^, Hyeju Oh^27^, Kousuke Onoue^67^, Hyojung Paik^44^, Wenchu Pan^68^, Bogyu Park^57^, Balint Armin Pataki^4^, Sumeet Patiyal^34^, Jian Peng^62^, Dimitri Perrin^24^, Kaike Ping^69^, Alidivinas Prusokas^70^, Augustinas Prusokas^71^, Peng Qiu^72^, Gajendra P.S. Raghava^34^, Derek Reiman^30^, Renata Retkute^73^, Roberto Romero^1,5-9^, Nay Min Min Thaw Saw^38^, Neelam Sharma^34^, Alok Sharma^74-77^, Ronesh Sharma^54^, Rahul Siddharthan^55^, Musalula Sinkala^78,79^, Marina Sirota^10,11^, Alex Soupir^32^, Marija Stanojevic^66^, Gustavo Stolovitzky^19,20^, Yufeng Su^62^, Alexander M. Sutherland^80^, András Szilágyi^37^, Mehmet Tan^22^, Adi L. Tarca^1,2,3^, Nandor G. Than^37,81^, Buu Truong^31^, Edwin Vans^54,77^, Fangping Wan^58^, Rohan B.H. Williams^82^, Wendy S.W. Wong^83^, Jeong Woong^57^, Li Xiaomei^31^, Dongchan Yang^43^, Sanghoo Yoon^39^, Dakota York^32^, James Young^32^, Thomas Yu^15^, Wei Zhu^84^

^22^Department of Computer Engineering, TOBB University of Economics and Technology, Ankara, Turkey

^23^School of Mathematics and Statistics, University of Hyderabad, Hyderabad, India

^24^School of Computer Science, Queensland University of Technology, Brisbane, Australia

^25^Departments of Computer and Data Sciences, Case Western Reserve University, Cleveland, OH, USA

^26^Department of Pharmaceutical Science, College of Pharmacy, University of South Florida, Tampa, FL, USA

^27^Department of Statistics, Daegu University, Gyeongsan, Republic of Korea

^28^Department of Computer Science and Engineering, Korea University, Seoul, Republic of Korea

^29^Department of Computer Science and Engineering, Indraprastha Institute of Information Technology, New Delhi, India

^30^Department of Bioengineering, University of Illinois at Chicago, Chicago, USA

^31^UniSA STEM, University of South Australia, South Australia, Australia

^32^Department of Biology and Microbiology, College of Natural Science, South Dakota State University, Brookings, SD, USA

^33^Max-Planck Institute of Molecular Plant Physiology, Potsdam, Germany

^34^Department of Computational Biology, Indraprastha Institute of Information Technology, New Delhi, India

^35^School of Electrical Engineering, University in Belgrade, Belgrade, Serbia

^36^Faculty of Information Technology and Bionics, Pázmány Péter Catholic University, Budapest, Hungary

^37^Institute of Enzymology, Research Centre for Natural Sciences, Budapest, Hungary

^38^Singapore Centre for Environmental Life Sciences Engineering, Nanyang Technological University, Singapore

^39^Division of Mathematics and Big Data Science, Daegu University, Gyeongsan, Republic of Korea

^40^Research School of Finance, Actuarial Studies & Statistics, Australian National University, Canberra, Australia

^41^ZS Associates India Private Ltd., Bangalore, India

^42^Department of Statistics, Stanford University, Stanford, CA, USA

^43^Department of Bio and Brain engineering, Korea Advanced Institute of Science and Technology, Daejeon, Republic of Korea

^44^Korea Institute of Science and Technology Information, Center for Supercomputing Application, Daejeon, Republic of Korea

^45^Health Science Center, Peking University, Beijing, China

^46^Department of Computer Science and Engineering, Korea University, Interdisciplinary Graduate Program in Bioinformatics, Korea University, Seoul, Republic of Korea

^47^Department of Electrical Engineering, Veermata Jijabai Technological Institute, Mumbai, India

^48^Bioinformatics Centre, CSIR-Institute of Microbial Technology, Chandigarh, India

^49^Bioinformatics Program, Department of Biology, Temple University, Philadelphia, PA, USA

^50^Patient-Centered Clinical Research Coordinating Center, National Evidence-based Healthcare Collaborating Agency, Seoul, Republic of Korea

^51^Munich School of Management and Munich Center of Health Sciences, Ludwig-Maximilians-University Munich, Germany, Munich, Germany

^52^SymbioSeq LLC, Ashburn, VA, USA

^53^Faculty of Pharmacy, University of Belgrade, Belgrade, Serbia

^54^School of Electrical and Electronic Engineering, Fiji National University, Suva, Fiji

^55^The Institute of Mathematical Sciences (HBNI), Chennai, India

^56^Institute of Health Economics and Health Care Management, Helmholtz Zentrum München, Munich, Germany

^57^DoAI Inc., Republic of Korea

^58^Institute for Interdisciplinary Information Sciences, Tsinghua University, Beijing, China

^59^Cancer & Stem Cell Biology Program, Duke-NUS Medical School, Singapore

^60^SingHealth Duke-NUS Institute of Precision Medicine, Singapore

^61^GeneDx, Gaithersburg, MD, USA

^62^Department of Computer Science, University of Illinois at Urbana-Champaign, Urbana, IL, USA

^63^ZS Associates International, Inc., London, UK

^64^CSIR National Chemical Laboratory, Pune, India

^65^Applied Artificial Intelligence Institute (A2I2), Deakin University, Victoria, Australia

^66^Center for Data Analytics and Biomedical Informatics, Computer and Information Sciences Department, Temple University, Philadelphia, PA, USA

^67^NEC Corporation, Tokyo, Japan

^68^Department of Statistics, Peking University, Beijing, China

^69^Guizhou Center for Disease Control and Prevention, Guiyang, Guizhou, China

^70^Plant and Microbial Sciences, School of Natural and Environmental Sciences, University of Newcastle, Newcastle, UK

^71^School of Life Sciences, Imperial College London, London, UK

^72^Department of Biomedical Engineering, Georgia Institute of Technology and Emory University, Atlanta, GA, USA

^73^Epidemiology and modelling group, Department of Plant Sciences, University of Cambridge, Cambridge, UK

^74^Department of Medical Science Mathematics, Medical Research Institute, Tokyo Medical and Dental University, Tokyo, Japan

^75^Institute for Integrative and Intelligent Systems, Griffith University, Brisbane, Australia

^76^Laboratory for Medical Science Mathematics, RIKEN, Yokohama, Japan

^77^School of Engineering and Physics, University of the South Pacific, Suva, Fiji

^78^University of Cape Town, Faculty of Health Sciences, Department of Integrative Biomedical Sciences, Computational Biology Division, Cape Town, South Africa

^79^University of Zambia, School of Health Sciences, Department of Biomedical Sciences, Lusaka, Zambia

^80^San Francisco, CA, USA

^81^Maternity Clinic of Obstetrics and Gynecology, Budapest, Hungary

^82^Singapore Centre for Environmental Life Sciences Engineering, National University of Singapore, Singapore

^83^Coherent Logic Limited, McLean, VA, USA

^84^CodexSage, LLC, Gaithersburg, MD, USA

