## Supplemental Figures and Table legends for "Crowdsourcing assessment of maternal blood multi-omics for predicting gestational age and preterm birth"

**Supplementary figures and tables**

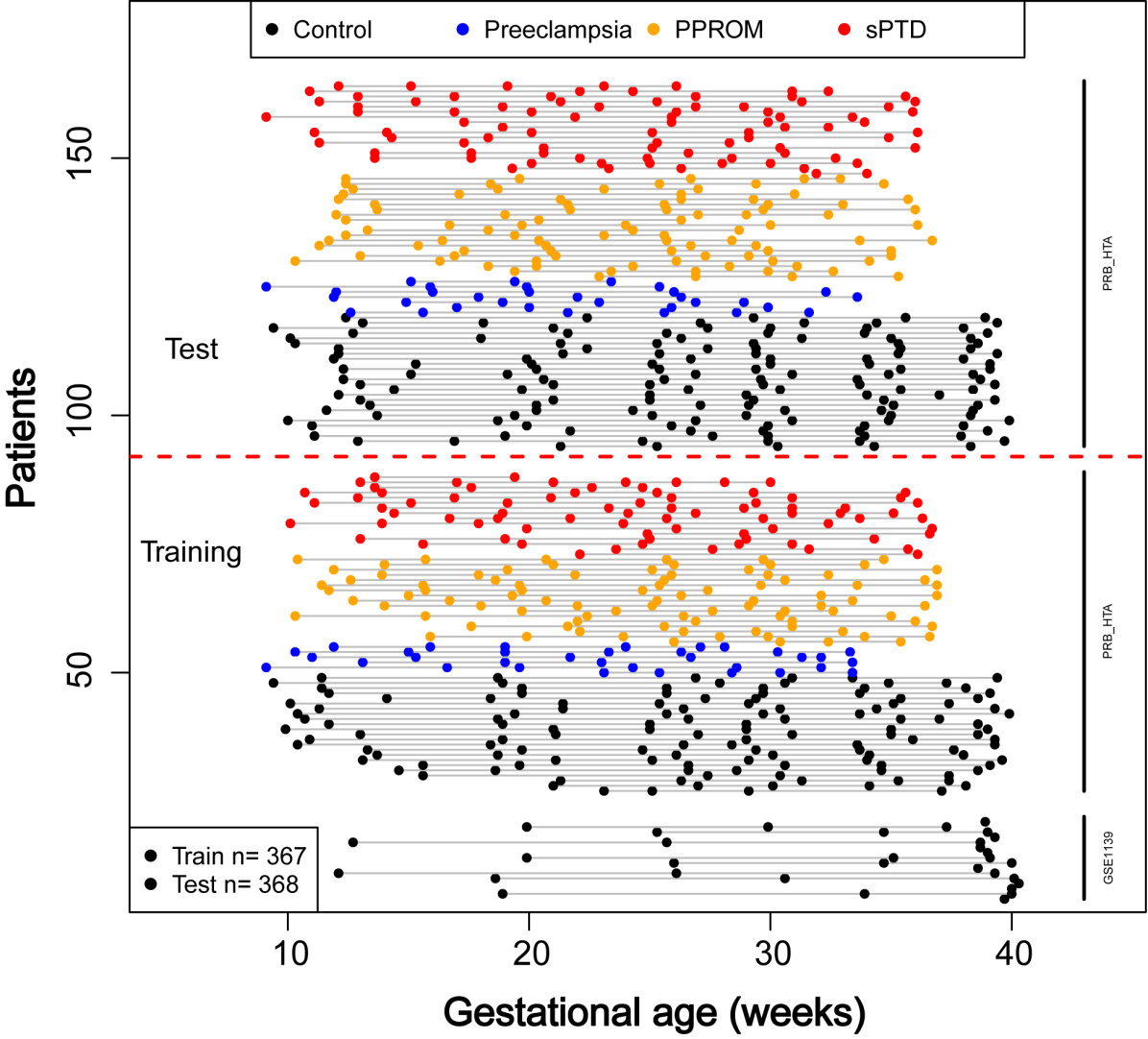

**Figure S1. Transcriptomics training and test set for sub-challenge 1.** GSE1139 is the GEO identifier of a previously released dataset while PRB_HTA refers to the transcriptomics dataset in **Fig 2**. Both datasets were generated using Affymetrix Human Transcriptome Arrays 2.0.

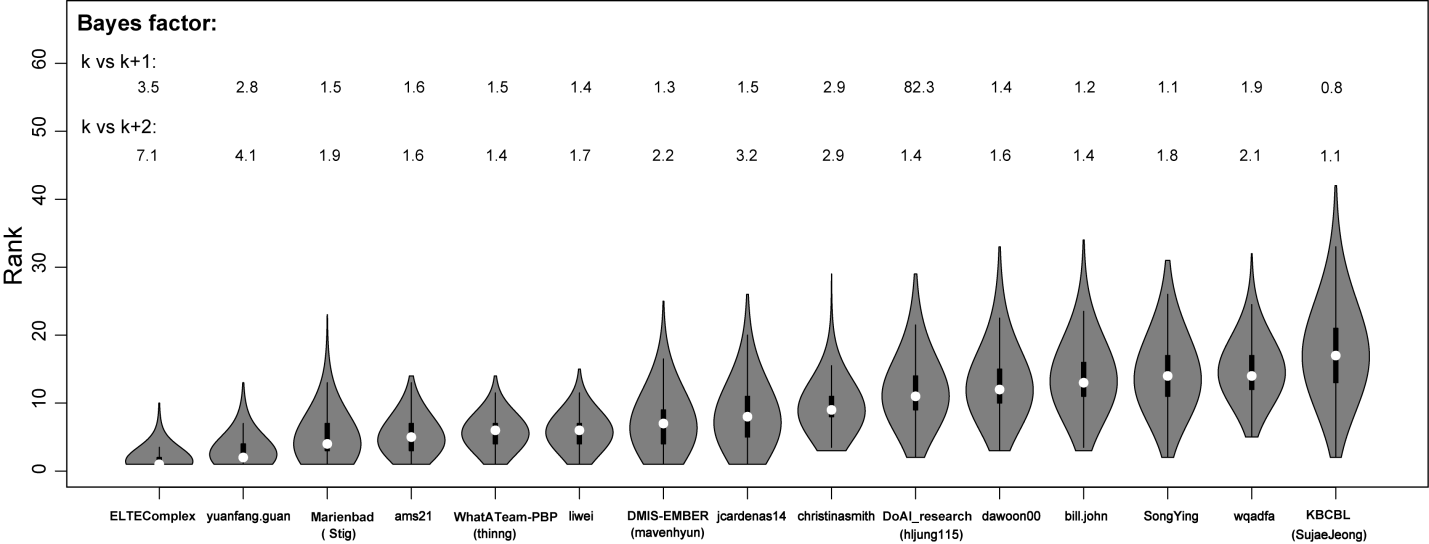

**Figure S2. Team rank stability in sub-challenge 1**. The violin plots show the distribution of the team ranks (the smaller the better) under bootstrap resampling scenarios of the test set. Bayes factors shown contrast the ranks of a given team (k) relative to the next ones (k+1 or k+2) defined by the official ranking.

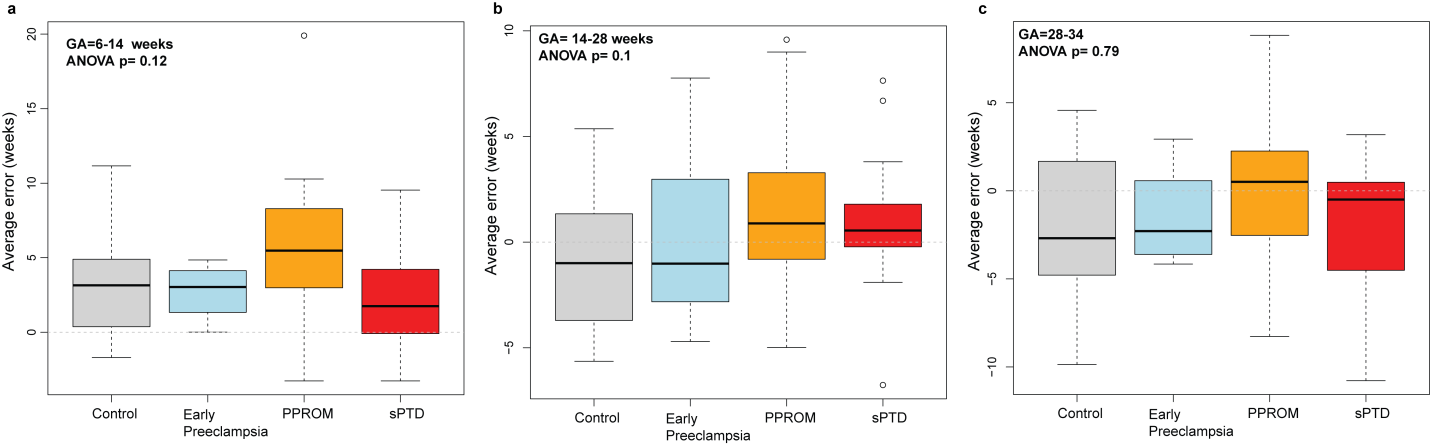

**Figure S3. Gestational age prediction error by pregnancy outcome and gestational age interval**. Test set prediction errors of model M_GA_Team1 were averaged over the eventual multiple samples of a given patient in the gestational age (GA) interval specified in each panel (A: 6-14, B: 14-28, C: 28-34 weeks). Note that early preeclampsia samples do not extend after 34 weeks. sPTD: spontaneous preterm delivery with intact membranes; PPROM: Preterm premature rupture of membranes.

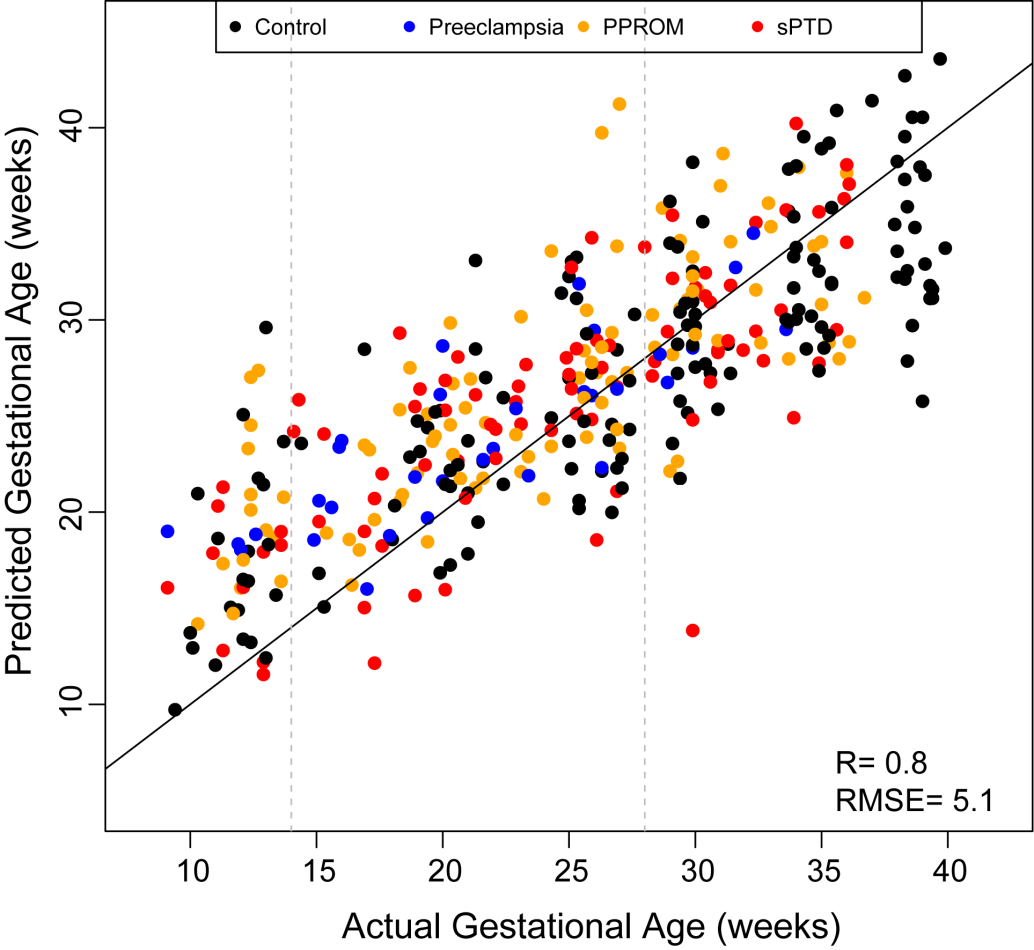

**Figure S4. Test set prediction of gestational age at blood draw by a core transcriptome model M_GA_Core**. R: Pearson correlation coefficient; RMSE: Root Mean Squared Error

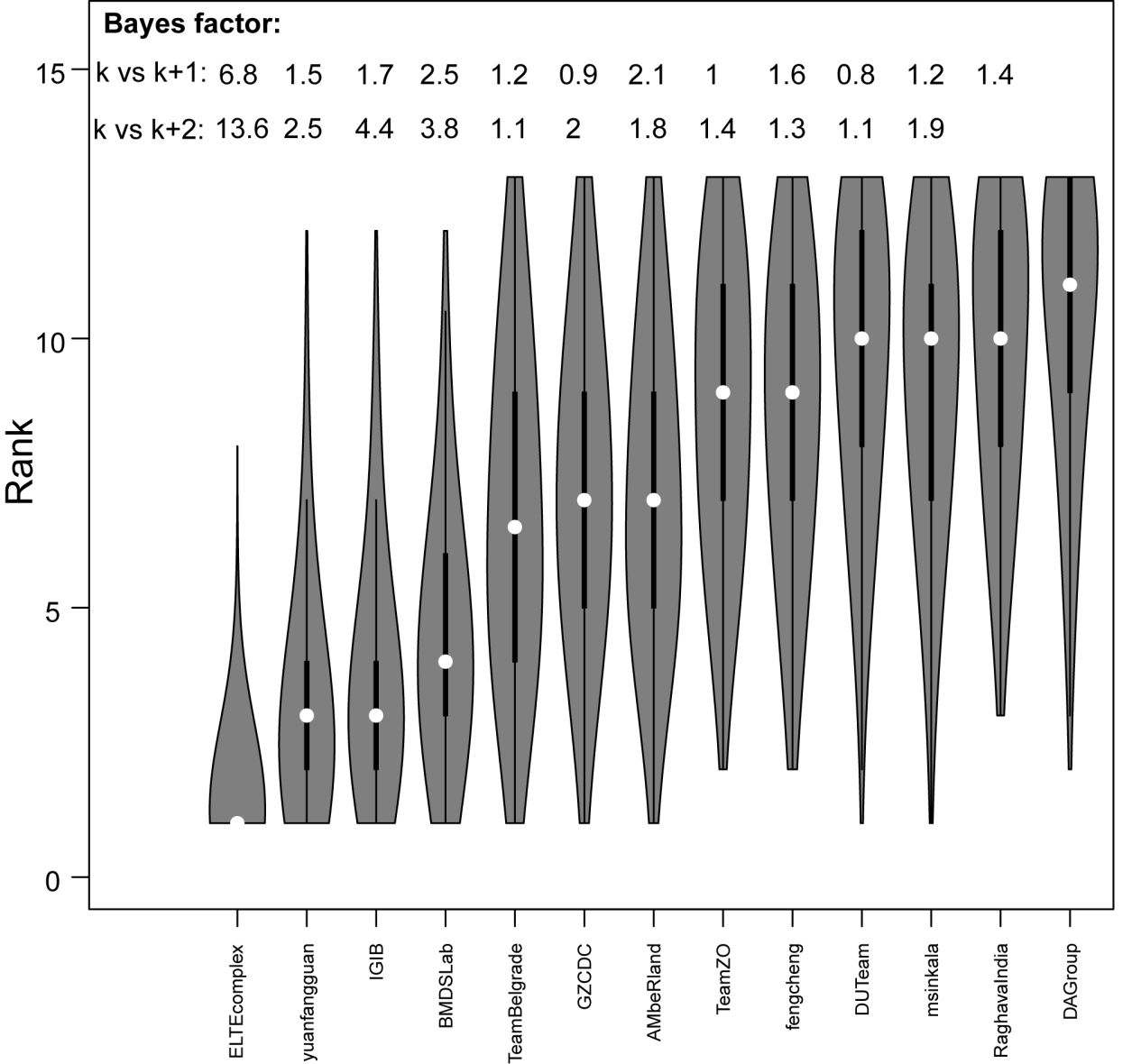

**Figure S5. Team rank stability analysis in sub-challenge 2**. The violin plots show the distribution of the team ranks (the smaller the better) under bootstrap resampling of the 10 train/test instances pertaining to each scenario, and also of the prediction criteria (columns in heatmap of **Fig. 8**). Bayes factors shown contrast the ranks of a given team (k) relative to the next ones (k+1 or k+2) defined by the official ranking.

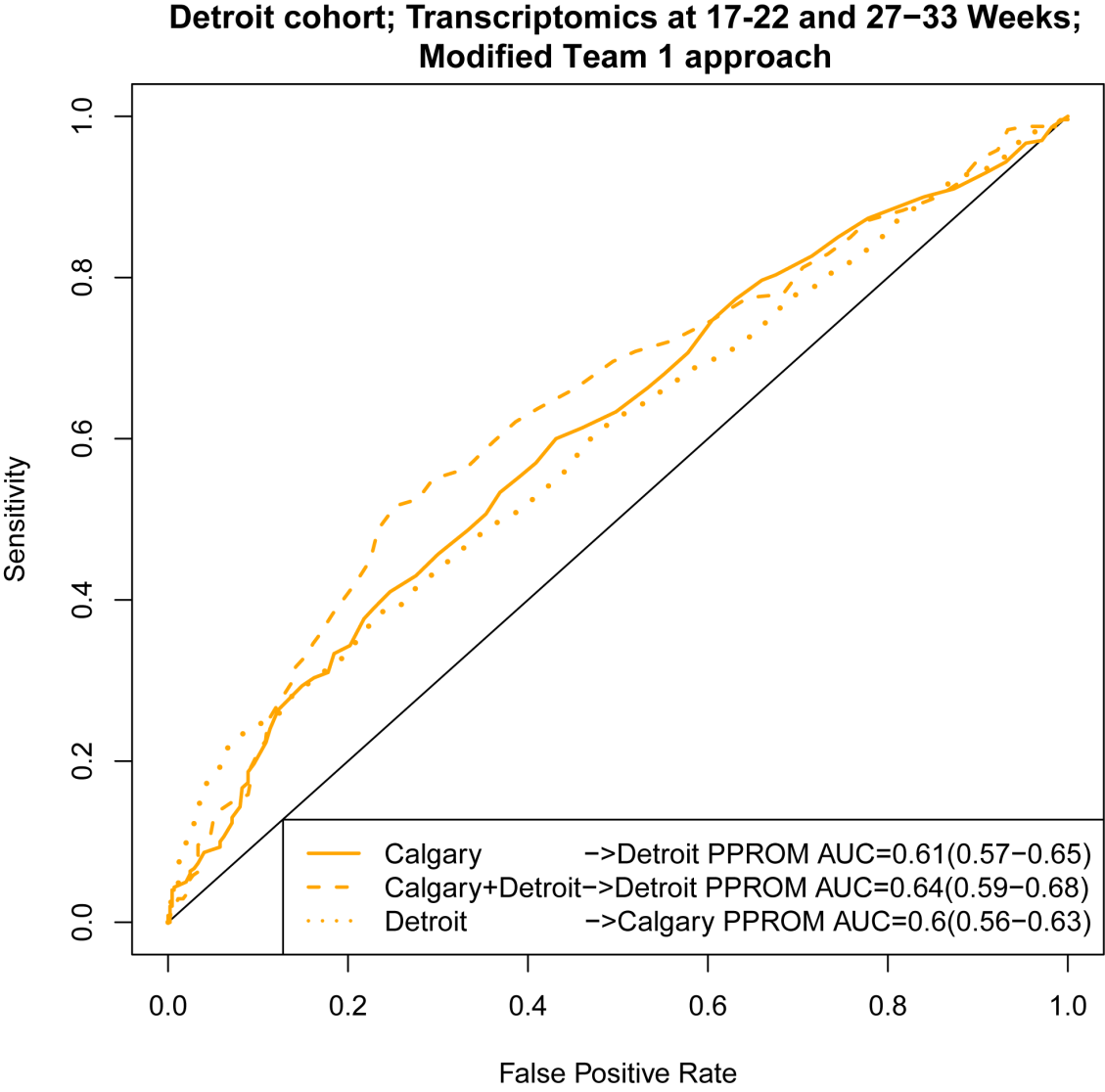

**Figure S6. Receiver operating characteristic curve (ROC) for prediction of PPROM across cohorts and microarray platforms using a modified approach of Team 1**. The original Team 1 approach selected genes by random forest model importance using data at the last available time point (27-33 weeks). The modified approach keeps all modeling aspects and the predictor genes the same, except that the gene expressions at the earlier time point (17-22 weeks) are also allowed as inputs in the random forest model.

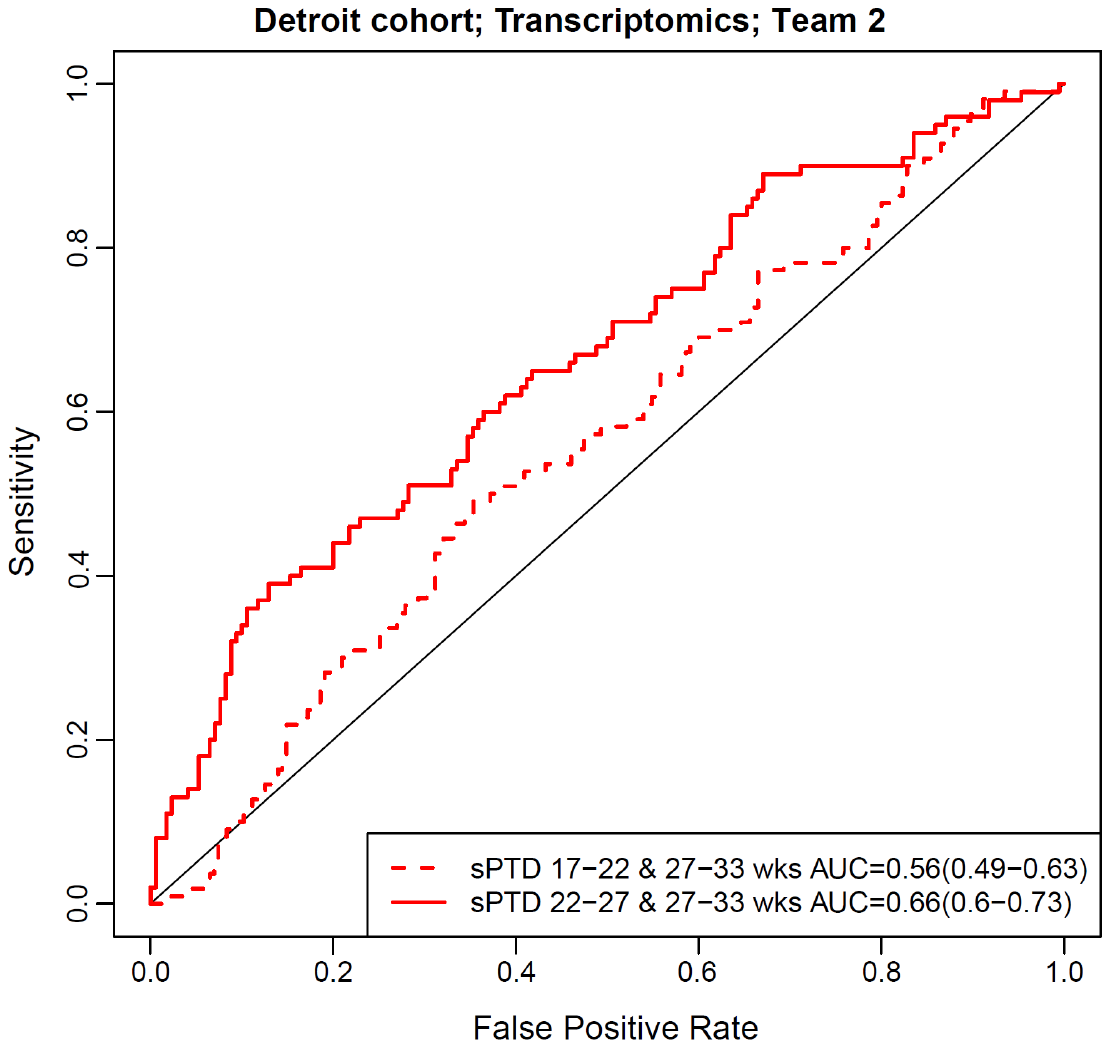

**Figure S7. Prediction of spontaneous preterm delivery by maternal whole blood transcriptomics in the Detroit cohort**. Receiver operating characteristic curve (ROC) represents the prediction of spontaneous preterm delivery by 50 genes using data from two different time points (17-22 and 27-33 weeks of gestation or 22−27 and 27−33 weeks of gestation) using the Team 2 approach.

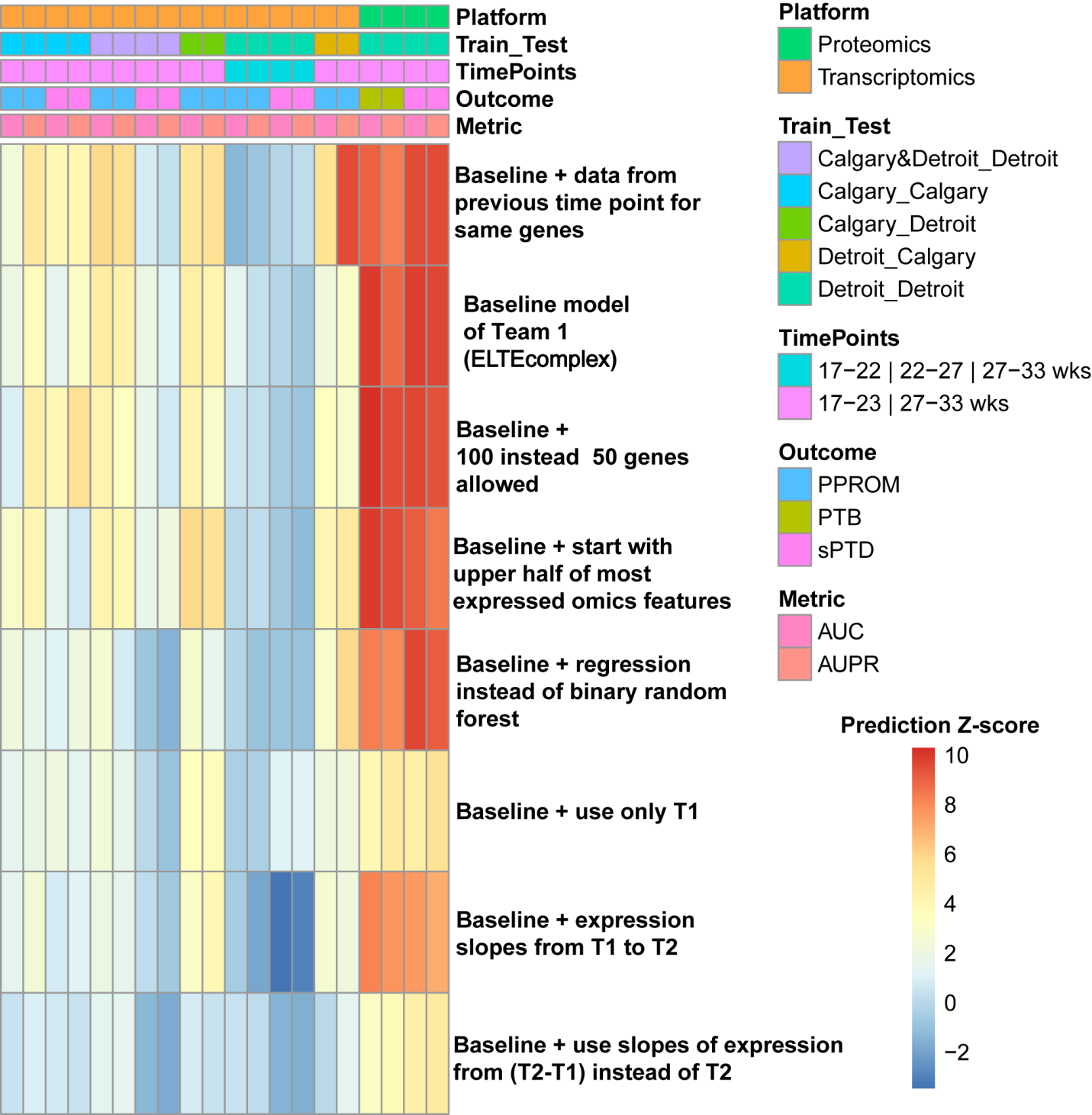

**Figure S8. Prediction performance of hybrid preterm birth prediction approaches by changes to the approach of Team 1**. Prediction performance is shown under the same scenarios shown in **Fig. 5**. AUROC and AUPRC metrics were converted into Z-scores and shown as a heatmap. The base line approach uses random forest model importance to select top 50 genes considering data at the last point (noted with T2 for both 2 time point and 3 time point scenarios) and then fits a random forest model with only those 50 genes. See result section for the meaning of the different alterations to the baseline approach shown as rows in the heatmap.

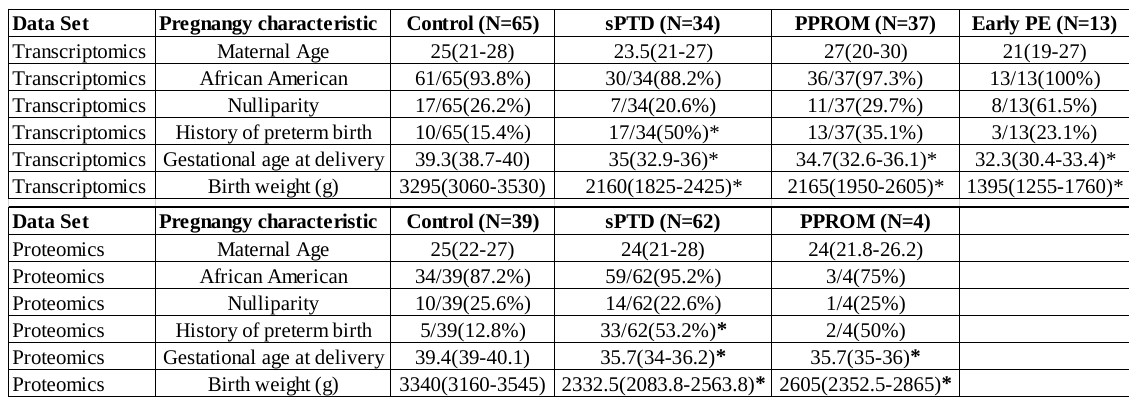

**Table S1. Demographic characteristics of the study population**. For each dataset, demographic and delivery information are presented as mean (inter quartile range) or number (%). Differences between disease groups and controls are assessed by t-test for continuous data or Fisher’s exact test for binary data, with * representing a significant difference (p<0.05).

**Table S2. Team prediction performance in sub-challenge 1**. The table shows the identifier of each of the 37 teams qualified in the final ranking of sub-challenge 1 as well as the best (smallest) Root Mean Squared Error (weeks) over up to 5 trials when predicting gestational age at delivery on the test set.

**Table S3. The 249-gene core transcriptome predicting gestational age M_GA_Core**. The gene symbol, ENTREZ database identifier and coefficient in the lasso regression model are shown.

**Table S4. Biological processes enriched among the genes part of the 249-gene core transcriptome predicting gestational age M_GA_Core**. A select group of biological processes enriched among these genes are shown using pie charts. GOBPID: Gene ontology biological process identifier; GOName: Biological process name; Count: Number of differentially expressed genes annotated to the biological process.	Size: Number of all genes on the microarrays annotated to the biological process. Odds Ratio: Enrichment analysis effect size; Pvalue: hypergeometric test p-value;	q: False Discovery Rate adjusted p-value.

**Table S5. Team prediction performance in sub-challenge 2**. For each combination of prediction scenario and outcome (see Table 1) the Z-score for both AUROC and AUPRC metrics are given for each team. After ranking teams by each of the 28 criteria, a sum of ranks and final rank were determined.

**Table S6. Gene changes preceding diagnosis with PPROM in Calgary and Detroit cohorts and supported by evidence from samples collected at 17-23 and 27-33 weeks**. Moderated t-test p-values and log2 fold changes (PPROM/Control) are included together with false discovery adjusted p-values.

**Table S7. Biological processes enriched in genes differentially expressed with PPROM**

**from Table S6.** A select group of biological processes enriched among these genes are shown using pie charts. GO Name: Gene ontology biological process name,	Count: Number of differentially expressed genes annotated to the biological process.	Size: Number of all genes on the microarrays annotated to the biological process.	Odds Ratio: Enrichment analysis effect size; Pvalue: hypergeometric test p-value;	q: False Discovery Rate adjusted p-value.

**Table S8. Gene expressions correlated with gestational age at delivery among controls and spontaneous preterm delivery cases in the Detroit cohort**. Correlation coefficients and p-values are given for each time point.

**Table S9. Plasma protein changes preceding diagnosis with sPTD the Detroit cohort at 27-33 weeks**. Moderated t-test p-values and log2 fold changes (sPTD/Control) are included together with false discovery adjusted p-values.

**Table S10. Plasma protein changes preceding diagnosis with sPTD the Detroit cohort at 17-22 weeks**. Moderated t-test p-values and log2 fold changes (sPTD/Control) are included together with false discovery adjusted p-values.
